# Site-specific processing of phosphoethanolamine cellulose by the BcsZ cellulase reveals stochastic biofilm cellulose modification

**DOI:** 10.64898/2026.08.24.745826

**Authors:** Jedrzej Rum, Jhih-Yi Huang, Elena N. Kitova, Theodore Tyrikos-Ergas, Ling Han, Martina Delbianco, John S. Klassen, Jochen Zimmer

**Author notes:** **Corresponding authors:** Jochen Zimmer, John S. Klassen **Email:** and.

## Abstract

Cellulose is a common component of bacterial biofilms where it interacts with other biopolymers to form a 3-dimensional matrix enclosing the bacteria. Synthesized and secreted by the synthase-dependent biosynthesis pathway common to many bacterial exopolysaccharides, its surface exposure depends on the presence of the periplasmic cellulase BcsZ. During export across the periplasm, *E. coli* and other *Enterobacteriaceae* modify cellulose with lipid-derived phosphoethanolamine (pEtN). How BcsZ hydrolyzes pEtN-cellulose in the periplasm is unknown and so is the native distribution pattern of pEtN on cellulose. Here, we used carbohydrate synthesis, X-ray crystallography, native mass spectrometry, and super-resolution MINFLUX nanoscopy to delineate BcsZ’s role during cellulose biosynthesis.

Crystal structures of BcsZ bound to chemically synthesized pEtN cello-oligosaccharides identify how the enzyme recognizes pEtN-modified glucosyl units. Comparing mono and double substituted cellohexaoses, we identify varying binding poses that are determined by two pEtN coordination sites within BcsZ’s catalytic pocket. Combined, our structural analyses reveal an ideal BcsZ cellohexaose ligand containing two pEtN modified units separated by an unmodified cellotriosyl unit. The enzyme binds and hydrolyzes this compound with substantially increased affinity and efficiency. Further, BcsZ digestion of native pEtN cellulose combined with native mass spectrometry analyses reveals the stochastic distribution of pEtN on biofilm cellulose. Additionally, MINFLUX co-localization of BcsZ with other components of the biosynthetic complex demonstrates BcsZ’s random distribution across the periplasm. Our data suggest BcsZ functions independently of the biosynthetic complex to clear mislocalized pEtN cellulose from the periplasm.

**Significance Statement:** Biofilms are an abundant form of bacterial growth and responsible for the majority of hospital-derived infections. Uropathogenic *E. coli* produces phosphoethanolamine cellulose as a stabilizing extracellular polysaccharide. Surprisingly, the periplasmic cellulase BcsZ, encoded in the cellulose biosynthesis operon, is necessary for efficient bacterial cellulose production. Crystal structures of BcsZ bound to chemically synthesized phosphoethanolamine cellulose fragments reveal how the enzyme recognizes and cleaves its unique substrate. Further, super-resolution fluorescence microscopy shows that BcsZ does not form a stable complex with other cellulose synthase components and likely diffuses in the *E. coli* periplasm. Finally, mass spectrometry of oligosaccharides released by BcsZ from biofilm *E. coli* indicates the stochastic modification of cellulose with phosphoethanolamine groups *in vivo*.

## Introduction

Biofilms, accounting for many nosocomial infections (1), are a common bacterial growth phase in which cells are surrounded by a 3-dimensional meshwork of biopolymers, including polysaccharides (2–5). The biofilm matrix (ECM) affects solute diffusion, intercellular communication, as well as the metabolic state of some bacterial sub-populations (6).

Cellulose is a common polysaccharide of enterobacterial biofilms (7). It is a linear polymer of glucosyl units linked via β-1,4 glycosidic linkages. The polysaccharide’s end carrying an unmodified C1 hydroxyl group represents the reducing end, while the opposing terminus is the ‘non-reducing end’. Cellulose is amphipathic, enabling it to self-assemble into fibrils or form composite materials with other carbohydrate- and non-carbohydrate-based biopolymers (8–10).

In Gram-negative bacteria, the cellulose synthase BcsA partners with additional non-enzymatic components to enable cellulose secretion across the cell envelope (11, 12). These include the periplasmic BcsB and outer membrane-integrated BcsC subunits. BcsA, BcsB and BcsC form the minimal machinery necessary for cellulose synthesis and secretion.

*Enterobacteriaceae* modify about half of cellulose’s glucosyl units with lipid-derived phosphoethanolamine (pEtN) to promote biofilm cohesion and resistance to sheer stress (13, 14). Modification occurs in the periplasm and is catalyzed by the membrane-embedded pEtN transferase BcsG (15–17).

An additional conserved subunit of bacterial cellulose biosynthetic systems is BcsZ, a family-8 glycosyl hydrolase that cleaves the β-(1,4)-linkages of cellulose. BcsZ is a periplasmic enzyme and, counterintuitively, is necessary for efficient cellulose secretion *in vivo*, in both cellulose- and pEtN cellulose-producing bacteria (16, 18). BcsZ exhibits a deep catalytic pocket that binds cello-oligosaccharides in a bent conformation (18, 20). The pocket is lined with aromatic and polar residues that coordinate cellulose through CH-π stacking and polar interactions at three binding sites before (−3 to −1) and two after (+1 to +2) the catalytic residues. These are a pair of acidic amino acids (Glu55 and Asp243 in *E. coli* BcsZ) that facilitate the S_N_2 nucleophilic attack of the nucleophilic water on the linkage connecting glucosyl units at subsites −1 and +1.

Previous cello-oligosaccharide-bound structures of BcsZ revealed how the enzyme recognizes unmodified cellulose (20). However, we currently lack any information on how BcsZ processes its native substrate, pEtN cellulose. Here, we present crystal structures of *E. coli* BcsZ bound to unmodified and pEtN-modified cellohexaose substrates. Our data reveals that BcsZ can accommodate pEtN groups at its −3 and +2 subsites, thereby dictating the oligosaccharide binding pose. Monosubstituted cellohexaoses bind such that the pEtN modified units occupy the preferred binding sites. Ligands carrying multiple pEtN units are accepted only when modified glucosyl units are separated by at least three unmodified units. Furthermore, BcsZ digestion of authentic *E. coli* pEtN cellulose reveals stochastic pEtN decorations, ranging from no to sparse and extensive modifications. MINFLUX nanoscopy (21) reveals that BcsZ does not co-localize with BcsA *in vivo*, consistent with a diffusion-based housekeeping function in the periplasm.

## Results

### Structural insights into BcsZ substrate coordination

To gain structural insights into BcsZ’s substrate coordination, the catalytically inactive enzyme (E55Q) (20) was crystallized either in the absence of ligands or the presence of cellopentaose or - hexaose by the sitting drop vapor diffusion method (see Methods), Table S1. Co-crystallization occurred in the presence of unmodified cellohexaose and –pentaose ligands, referred to as AAAAAA and AAAAA, with A denoting an unmodified glucosyl unit. The apo crystals were soaked with synthesized pEtN cellohexaoses (22, 23). This ensured an almost uniform ligand occupancy of the catalytic pockets of the four BcsZ protomers in the asymmetric unit, Fig. S1A. The ligands soaked were PAAAAA, AAAAAP, AAPPAA, and APAAPA. Here, ‘P’ denotes a pEtN-derivatized glucosyl unit and the position within the string (left to right, non-reducing to reducing end) indicates its distance from the non-reducing end sugar.

### BcsZ binds cellulose in a 3+2 binding pose

Comparing how cellohexaose and cellopentaose bind to BcsZ reveals a shared coordination of five glucosyl units within the catalytic pocket, Fig. 1A-C. The ligands are positioned such that the reducing terminal disaccharide unit occupies subsites +1 and +2 past the catalytic residues Glu55(Q) and Asp243. At these sites, Tyr332 and Tyr242 form CH-π stacking interactions with the sugars and Asp243 hydrogen bonds with the C6 hydroxyl of the glucose moiety at position +1, Fig. 1B and C. Both monosaccharide units adopt the energetically favored chair conformation ^4^C_1_. The next glucosyl unit at position −1, connected to the preceding disaccharide via the scissile bond, is in a boat-like conformation and coordinated via its C2 and C3 hydroxyls and the C4 oxygen by Arg246, Asp116, and Tyr182, respectively. The putative nucleophilic water is positioned ~3.3 Å above the C1 carbon. It is coordinated by hydrogen bonds involving Asp243, Tyr182, the C6 hydroxyl group of the preceding glucosyl unit at subsite +1, as well as an additional water molecule above the ring oxygen of the same glucosyl unit, Fig. 1D, S1B, as observed in the GH8 cellulase from *Clostridium thermocellum* (24, 25). The following two glucosyl units at subsites −2 and −3 are again in the energetically favored ^4^C_1_ chair conformation. At subsite −2, the sugar stacks against Trp96 and its C6 hydroxyl group hydrogen bonds with the backbone carbonyl of Ser113. Subsite −3 lacks any characteristic stacking interactions. Instead, the glucosyl moiety is held by polar interactions between its C2 hydroxyl group with the sidechains of Asp110 and Asn112, Fig. 1C.

**Fig. 1.**
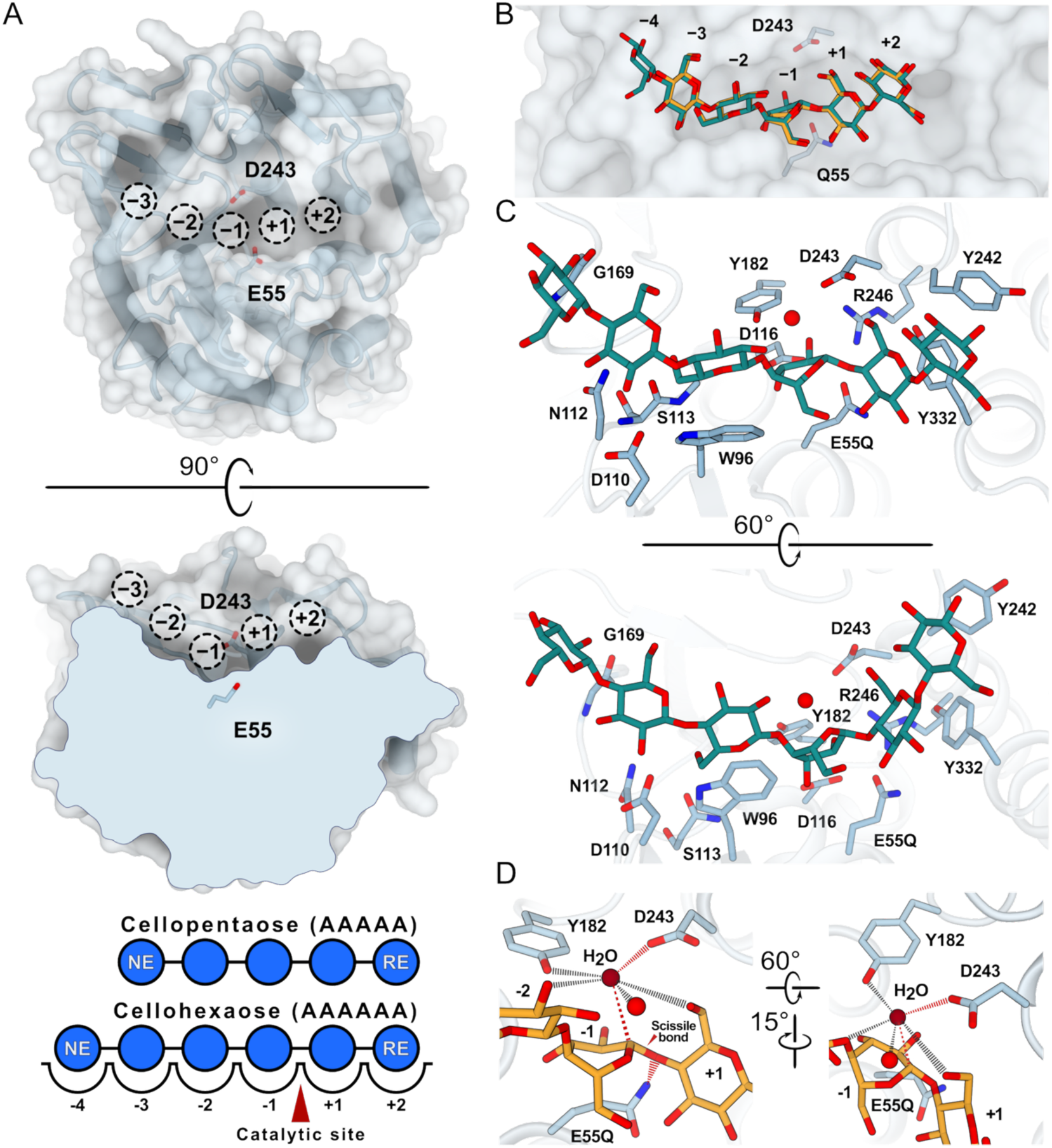
Structure of cellopentaose and cellohexaose bound BcsZ. (A) Cartoon and surface representation of BcsZ (pdb 3QXQ) highlighting the catalytic residues and glucosyl binding sites. Bottom: Illustration of the catalytic pocket with glucosyl units shown as blue circles. RE/NE: reducing and non-reducing ends. (B) Superimposition of cellopentaose (orange) and cellohexaose (teal blue) in the binding pocket. (C) Details of cellohexaose coordination. (D) Close-up view of the active site. The catalytic water is shown as a dark red sphere. The path for the nucleophilic attack on the C1 carbon is shown as a red dashed line. Hydrogen bonds are indicated as black dashed lines.

In the cellohexaose-bound structure, the coordination of the oligosaccharide is unchanged, with the exception that the additional glucosyl unit extends past subsite −3 at the non-reducing end, Fig. 1C. The only noticeable interaction of this sugar unit with BcsZ occurs via a hydrogen bond between its C2 hydroxyl group and the backbone carbonyl oxygen of Gly169. We refer to this site as subsite −4.

Of the four protomers in the asymmetric unit, two show continuous electron density for the ligand over the scissile bond. In the other two cases, however, the connecting oxygen between the glucosyl units at subsite −1 and +1 is weak, if not missing at a contour level that resolves the rest of the ligand, Fig. S1B. However, even in this case, the nucleophilic water is well resolved in all cases, suggesting that the ligands are not hydrolyzed but the glycosidic bond may be flexible.

### Position dependent recognition of pEtN cello-oligosaccharides

We next tested whether cellohexaose ligands with pEtN modifications at the reducing or non-reducing ends (AAAAAP and PAAAAA, respectively) can bind to BcsZ’s catalytic cleft, Fig. 2 A-D. Indeed, the AAAAAP ligand binding pose is superimposable with the cellohexaose position described above, Fig. 1B, 2B, C. The pEtN-modified unit at subsite +2 stacks against Tyr242 and the pEtN phosphate moiety attached to its C6 hydroxyl fits into a pocket underneath the N-terminus of BcsZ’s last α-helix (residues Ala330-Trp343). In this position, the phosphate group is in hydrogen bonding distance to the backbone amide groups of Tyr331, Tyr332 and Asn333, situated in the first helical turn. Mediated by water molecules, the phosphate moiety also interacts with Asp44 as well as the side chain of Asn333. Further, the terminal amino group of the ethanolamine moiety is stabilized via hydrogen bonds with Ser46 and the backbone carbonyl of Asp329.

**Fig. 2.**
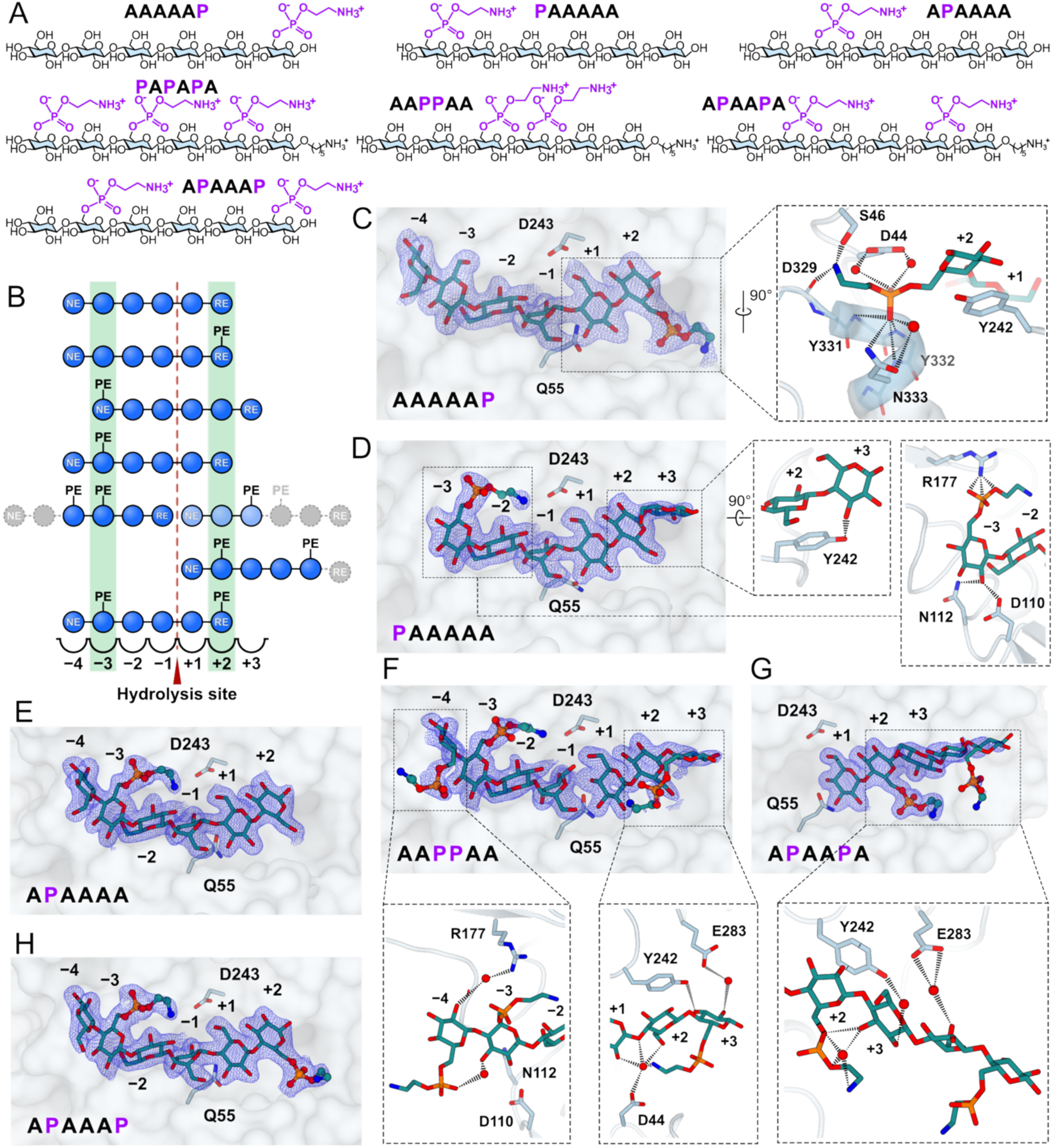
Binding of phosphoethanolamine cellulose to BcsZ. (A) Chemical structures of the pEtN modified cellohexaoses used in this study. ‘A’ and ‘P’ denote unmodified and pEtN modified glucosyl units, respectively. (B) Cartoon illustration of the binding poses observed for the various ligands. The –3 and +2 coordination sites are highlighted. Grey symbols indicate glucosyl units that could not be resolved in crystal structures. (C and D) 2FoFc electron densities of the terminally modified ligands in BcsZ’s catalytic pocket and close-up view on the coordination. The density is contoured at 1σ. (E-H) As for panels C and D for the indicated ligands.

When the pEtN group is located at the non-reducing end of the cellohexaose ligand (PAAAAA), the molecule binds in a new pose with three glucosyl units at the positive and negative subsites each, Fig. 2D B, and S2. The ‘P’ moiety is positioned at subsite −3. Here, the pEtN group is coordinated by Arg177, which is conserved among BcsZ’s from pEtN cellulose-producing bacteria, Fig. S3. Its guanidinium group is in hydrogen bonding distance to the ester bond linking pEtN’s phosphate and ethanolamine moieties. A second, likely weaker, interaction exists between the phosphate moiety, the Arg177 guanidinium group, a mediating water molecule, and the backbone of Gly169.

In the PAAAAA binding pose, the reducing end moiety extends past subsite +2, thereby identifying a +3 binding site, Fig. 2D. At this site, the sugar only forms a single interaction with BcsZ, between its C3 hydroxyl group and the side chain of Tyr242. It is further stabilized by a hydrogen bond between its C6 hydroxyl group and the C2 hydroxyl of the preceding glucosyl unit.

If the ‘P’ group resides at the second cellohexaose position (APAAAA), however, the ligand is bound as observed for the unmodified cellohexaose and AAAAAP ligands, Fig. 2E, B, S2. In this case, the ‘P’ moiety also resides at subsite −3 and mediates the same interactions as described above for the register shifted PAAAAA ligand, including the interaction with Arg177. This suggests that the pEtN interaction with Arg177 likely determines the ligand’s overall binding pose.

### Varying binding poses for multi-substituted cellohexaose ligands

To probe the acceptance of pEtN groups at other positions, we designed three additional pEtN cellohexaoses, two harboring two and one containing three pEtN modifications, Fig. 2A. Crystallographically, we failed to detect binding of the triple modified construct (PAPAPA).

The double modified ligands were AAPPAA and APAAPA, Fig. 2A. Our crystal structures reveal that ligand AAPPAA binds BcsZ in two alternative poses, Fig. 2B, F. The first pose positions the (AA)<u>PPAA</u> portion of the ligand (denoted ‘reducing end segment’) at subsites −4 through −1, the other places the <u>AAP</u>(PAA) trisaccharide (denoted ‘non-reducing end segment’) at subsites +1 and +3 and beyond.

The reducing end segment (AA)<u>PPAA</u> is bound such that the pEtN-modified glucosyl units are localized at subsite −4 and −3. The interactions at subsite −3 are as describe above for the PAAAAA ligand. However, in this case, the ‘P’ sugar is preceded by another pEtN-modified glucosyl unit whose pEtN moiety points roughly in the opposite direction, Fig. 2F. The only stabilizing interaction of this pEtN group appears to be a water-mediated hydrogen bond of its phosphate group with the C3 hydroxyl of the glucosyl unit at subsite −3. Accordingly, its ethanolamine moiety is poorly resolved. The ligand’s non-reducing end segment <u>AAP</u>(PAA) binds with both of its unmodified glucosyl units to subsites +1 and +2 and positions the pEtN modified sugar at subsite +3, Fig. 2B, F. Here, the C2 hydroxyl group of the P sugar interacts with Glu283, and its C3 hydroxyl contacts the side chain of Tyr242, as described above for the PAAAAA substrate. In addition, the attached pEtN group, and in particular its terminal amino group, is involved in a hydrogen bonding network including the oxygen atom of the glycosidic linkage connecting sugars at subsites +1 and +2, a water molecule, as well as Asp44. In contrast, the APAAPA ligand binds in a single binding pose but only to subsites +1 to +3 and beyond, Fig. 2B, G. At least four of its glucosyl units are resolved in all protomers of the asymmetric unit, with a pEtN-modified unit located at subsite +2. This suggests that the ligand can only bind with its non-reducing end segment <u>APAA</u>(PA) to BcsZ’s catalytic cleft. The pEtN-modified glucosyl unit is coordinated as described above for the AAAA<u>AP</u> ligand, with its phosphate moiety situated beneath the N-terminal end of BcsZ’s C-terminal α-helix, Fig. 2C. Additionally, the C3 hydroxyl group of the sugar unit at subsite +3 breaks the interaction with Tyr242 that was observed in the PAAAAA and AAP(PAA)-bound poses. Instead, it forms hydrogen bonds with the phosphate moiety of the pEtN group at subsite +2 and a water molecule, thereby ensuring co-planarity of the glucosyl units at subsites +2 and +3. Tyr242 now interacts with the C2 hydroxyl of the +3 sugar, bridged by a water molecule. The electron density of the fourth glucosyl unit resolved past subsite +3 is weak but discernible. Here, Glu283 interacts with C6 hydroxyl, via a water molecule, Fig. 2G and S1.

### The ideal ligand contains two pEtN groups separated by an unmodified cellotriosyl unit

Our crystallographic analyses suggest that BcsZ could accommodate two pEtN groups in its catalytic cleft at subsites −3 and +2, corresponding to an APAAAP ligand, Fig. 2A. Other modification patterns lead to alternative, perhaps product-mimicking, binding poses. To test this hypothesis, we synthesized the APAAAP ligand. Crystallographic analysis demonstrates that the ligand indeed binds to BcsZ over its entire catalytic cleft, with four glucosyl units residing at positions −4 to −1 and two at sites +1 and +2, Fig. 2B, H. As expected, the pEtN-modified sugars are at positions −3 and +2 and are coordinated as described above for the PAAAAA and AAAAAP-bound structures. No additional conformational changes of either BcsZ or the oligosaccharide are observed with this ‘ideal’ substrate.

### Substrate recognition and kinetics of pEtN cellulose hydrolysis

Native electrospray ionization mass spectrometry (nMS) was employed to determine the binding affinities of BcsZ for the synthesized pEtN cellohexaoses (26). By this method, the absolute ratio of ligand-bound and apo BcsZ at different BcsZ-to-ligand ratios can be determined, resulting in absolute dissociation constants (see Methods). To this end, the catalytically inactive BcsZ-E55Q mutant, used for our crystallographic studies, was incubated with the various ligands, Fig. 2A, and subjected to nMS.

BcsZ binds unmodified cellohexaose with a dissociation constant (K_D_) of about 0.36 mM, Table 1. This K_D_ is reduced 4-fold to 0.09 mM if the ligand carries pEtN groups at glucosyl units two and six (APAAAP), corresponding to the ‘ideal’ binder based on crystallography, Fig. 3A-C. In contrast, the K_D_ for the PAPAPA ligand not detected crystallographically is increased over 20-fold to 8 mM, Fig. 3B, perhaps due to partial binding to BcsZ’s catalytic cleft. Ligands harboring pEtN groups at either the −3 or +2 subsites of the catalytic pocket, APAAAA or AAAAAP, showed increased binding affinities relative to unmodified cellohexaose with K_D_s of 0.21 and 0.13 mM, respectively, Table 1 and S5. This suggests that additive effects account for the highest affinity binding of the APAAAP ligand. Coordination of pEtN at subsite +2 primarily accounts for higher affinity binding, indicated by both the lower K_D_ of ligand AAAAAP, compared to PAAAAA and APAAA, and the stronger binding of APAAPA, relative to AAPPAA, Table 1.

**Fig. 3.**
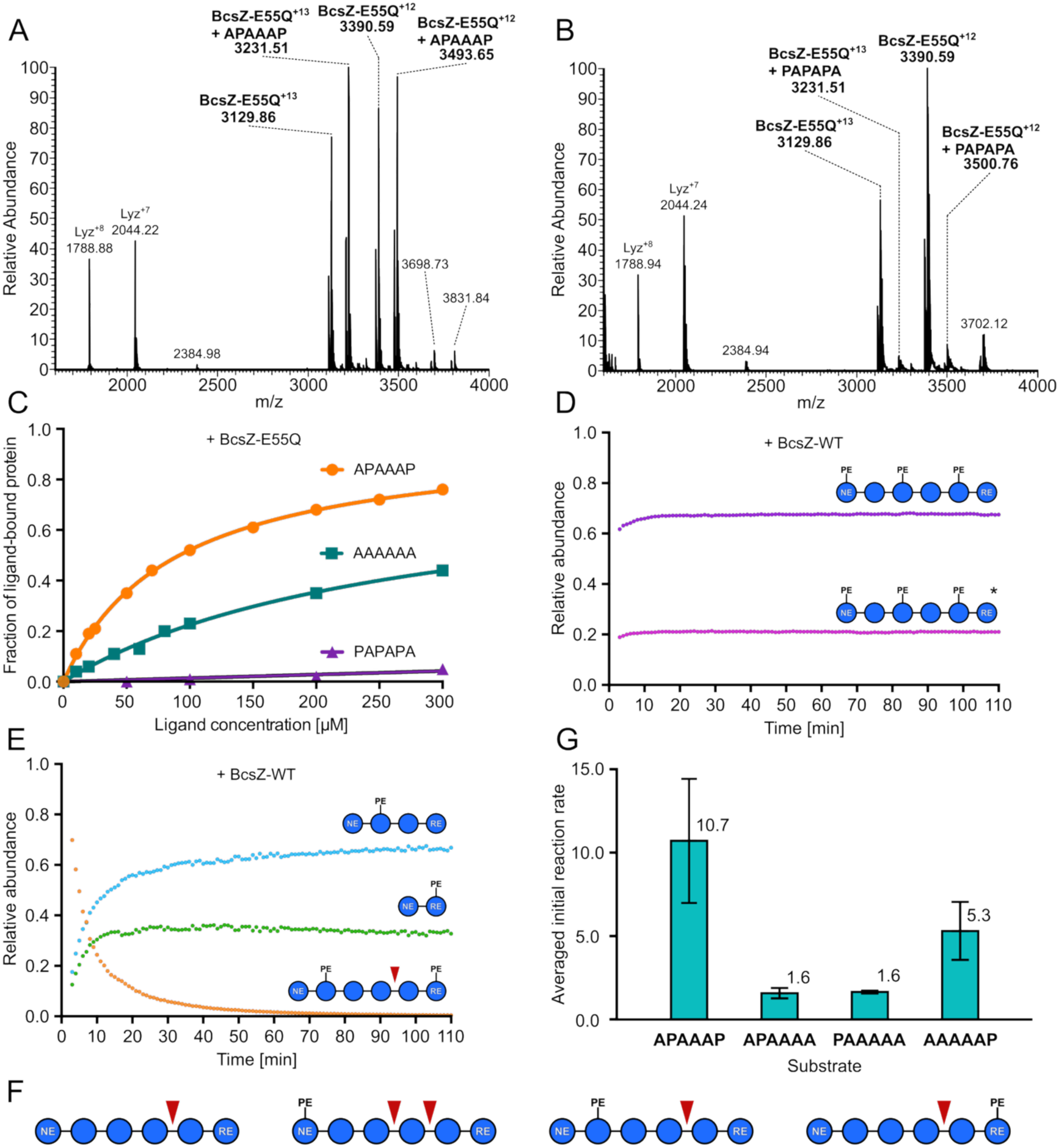
Native mass spectrometry analysis of pEtN cellulose binding and hydrolysis. (A and B) Mass spectrum of the catalytically inactive BcsZ-E55Q mutant bound to the ideal and weakest binders, respectively. (C) Quantification of ligand bound complexes as a function of ligand concentration. (D) Mass spectrometry detection of the weakest binder (PAPAPA) in the presence of wild type BcsZ over the indicated time interval. No hydrolysis product is observed. * denotes the doubly protonated species of the same substrate. (E) As for panel (D) but for the ideal binder. Hydrolytic fragments are detected within a few minutes. (G) Averaged initial reaction rates for turnover of the indicated ligands, relative to unmodified cellohexaose. (F) Cartoon representation of the observed cleavage products relative to the pEtN modification pattern.

**Table 1:**
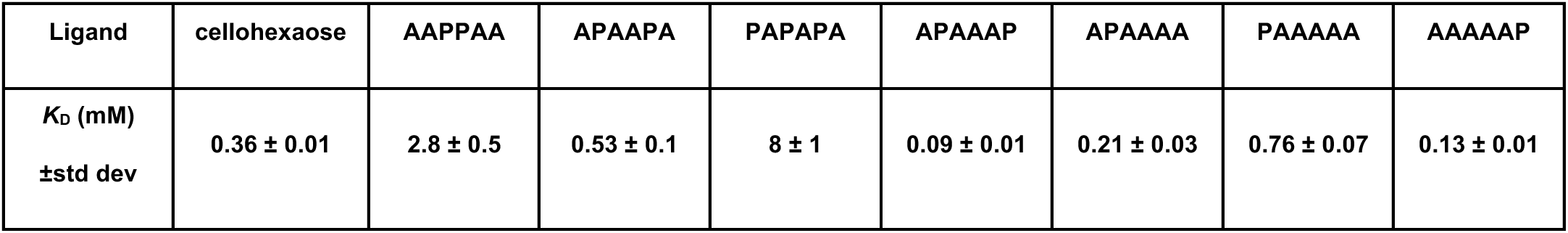

Native MS was also used to monitor the cleavage of the synthetic pEtN ligands. Wild-type BcsZ was incubated with each substrate for up to 120 min, during which the intact substrate and its cleavage products were continuously monitored by nMS (see Methods). As expected from its binding pose, BcsZ mostly cleaves unmodified cellohexaose into di- and tetrasaccharide fragments. Small amounts of cellotriose were also detected Fig. S6. No hydrolysis was observed for the AAPPAA, APAAPA, and PAPAPA compounds, which either did not bind at all to BcsZ or were positioned in alternative ‘product-mimicking’ binding poses in the crystal structures, Fig. 2F-G, 3D and S6.

The APAAAA substrate was cleaved into an unmodified cellobiose unit (AA) and cellotetraose carrying a pEtN modification at its second glucosyl unit from the non-reducing end (APAA), Fig 3F and S6. In contrast, the PAAAAA ligand produced two detectable cellotriose units, one unmodified, and the other containing a pEtN unit at the non-reducing end (AAA and PAA), as well as pEtN cellotetraose (PAAA) and cellobiose (AA) units, Fig. S2 and S6. The PAA and AAA products are in agreement with the preferred positioning of the ‘P’ group at subsite −3, Fig. 2D. The second product set (PAAA and AA) likely arises from an alternative binding pose, similar to the one observed for unmodified cellohexaose or the AAAAP ligand. Here, the ‘P’ moiety would occupy the −4 and the reducing end AA disaccharide the +1 and +2 subsites. Next, the cellohexaose ligand with a reducing end pEtN moiety (AAAAAP) was hydrolyzed into pEtN-modified cellobiose (AP) and unmodified cellotetraose (AAAA), Fig. S6, as predicted from the corresponding crystal structure, Fig. 2C. Lastly, the ideal double substituted cellohexaose ligand (APAAAP) was rapidly cleaved into pEtN-modified cellobiose and cellotetraose units (AP and APAA, respectively), Fig. 3E.

Comparing the initial hydrolysis rates of the pEtN-substituted substrates reveals position-dependent variability, Fig. 3G and S7. Relative to unmodified cellohexaose, the hydrolysis rate for the ideal APAAAP substrate was increased about 11-fold. The mono-substituted ligands, APAAAA and PAAAAA, were both hydrolyzed at roughly 2-fold higher rates while the AAAAAP ligand was degraded about 5-fold faster than the unmodified ligand. Hence, the pEtN-mediated increased hydrolysis rates are additive, especially when comparing turnover of the APAAAA and AAAAAP ligands, with the main contribution resulting from placing a ‘P’ moiety at position +2.

### Release of pEtN cello-oligosaccharides from native pEtN cellulose

Knowing BcsZ’s substrate preference, we utilized the enzyme to gain insight into the pEtN distribution pattern on native pEtN cellulose. To this end, pEtN cellulose was isolated from *E. coli* expressing the entire Bcs machinery, yielding cellulose indistinguishable from cellulosic material isolated from the reference strain UTI189, as previously described (15, 16). The isolated material was degraded by BcsZ, and the released oligosaccharides were characterized using nMS.

Among the various oligosaccharides detected, nMS confirmed the presence of unmodified units alongside pEtN cello-oligosaccharides containing two to nine glucosyl units, Fig. 4A-C, S8A, C, D and Table S2. In addition, mono substituted species of four and five glucosyl units alongside cello hexa- to octaoses with two to four pEtN modifications were identified, Fig. 4B-C and S8C. Similar oligosaccharides were also detected after exposing intact *E. coli* cells producing pEtN cellulose to BcsZ Fig. S8B, indicating that the identified fragments indeed represent surface exposed pEtN cellulose, Fig. S8B. Combined, this suggests varying degrees of cellulose pEtN modification *in vivo*, Fig. 4D.

**Fig. 4.**
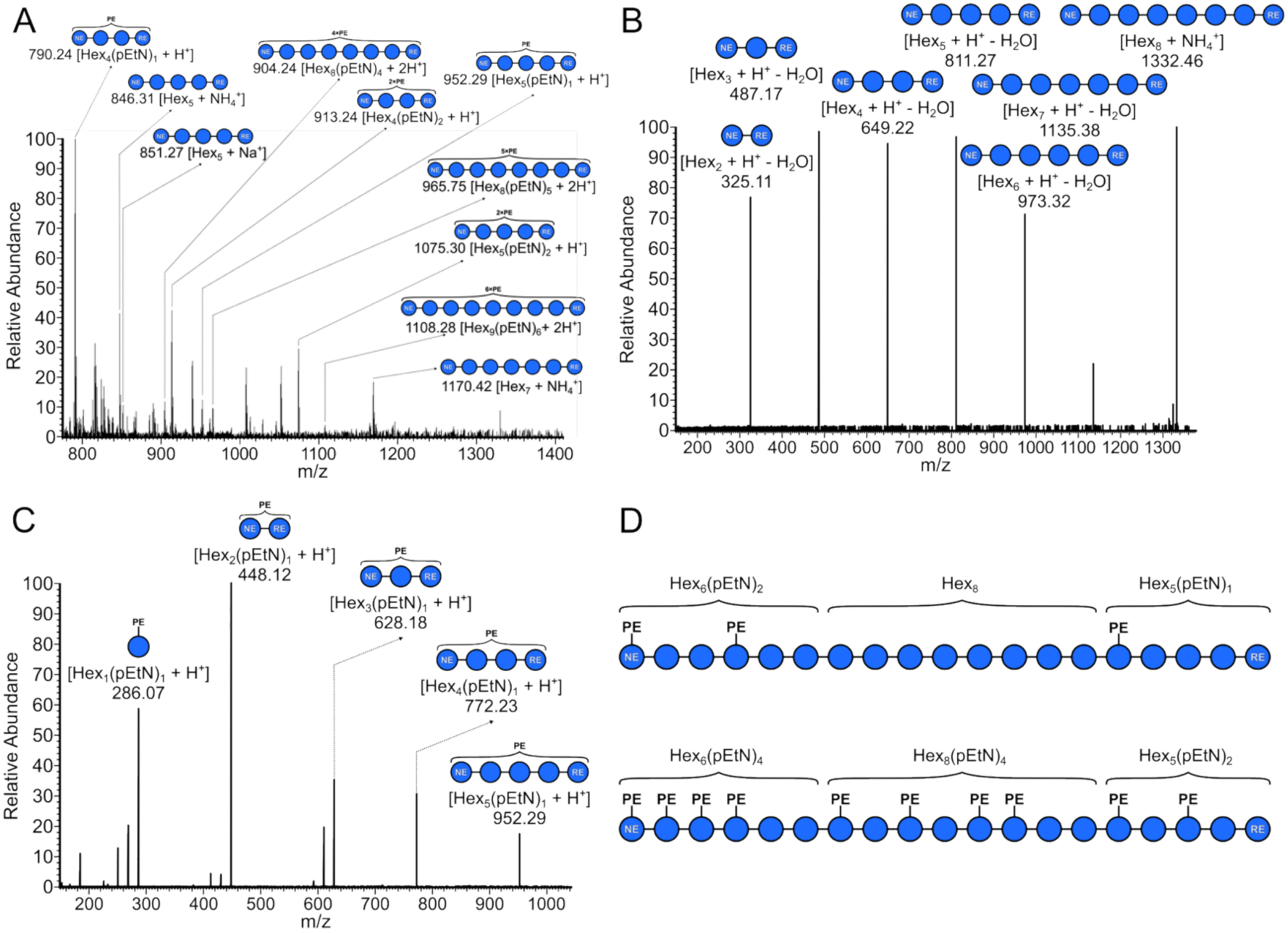
Mass spectrometry analysis of native pEtN cellulose fragments. (A) Cello-oligosaccharides released by BcsZ from in vivo produced pEtN cellulose, the mixture contains both modified and unmodified fragments. (B) Example of MS/MS spectrum confirming the identity of unmodified cellooctaose units released from purified pEtN cellulose. (C) Representative MS/MS spectrum of mono-substituted cellopentaose, released from purified pEtN cellulose. (D) A possible pEtN distribution pattern on E. coli produced cellulose based on the detected digestion products.

### In vivo localization of BcsZ

BcsZ may not need to stably interact with the cellulose biosynthetic Bcs machinery to degrade periplasmically mislocalized cellulose. To test its interaction with the inner membrane-integrated Bcs components, we employed MINFLUX nanoscopy (27) to co-localize fluorescently tagged BcsZ, alongside BcsA, the cellulose synthase. To this end, pairs of fluorescent markers were introduced into (a) BcsA (labeled with an AlexaFluor 647 conjugated antibody) and BcsZ (fused to Halo labeled with AlexaFluor 660), (b) BcsA and BcsB (labeled with Flux 680), and (c) BcsA and β-lactamase (antibody labeled with AlexaFluor 680), Fig. S9A. The emissions of the respective fluorophores can be assigned after data acquisition using a dual camera configuration in the cyanine (Cy5) near and far range, as described (28). Co-localization of BcsA and BcsB served as a positive control for Bcs components present in the same complex (15), while BcsA and β-lactamase are not expected to interact. The Halo-tagged BcsZ was confirmed to be periplasmically expressed and catalytically active by macro-colony staining, as described (16), Fig. S9B.

The obtained fluorophore localizations were processed for each pair based on nearest neighbor (NN) analysis (see Methods). In short, we analyzed how many BcsB-, BcsZ-, or β-lactamase-associated fluorophores co-localized with BcsA within a distance of 50 to 300 nm. Considering antibody labeling of BcsA, the upper distance limit of the NN analysis corresponds to about 10-times the largest fluorophore separation in the BcsA-BcsB complex (15).

MINFLUX localization of BcsA reveals a sparse but roughly uniform localization at the cell periphery, consistent with its integration into the inner *E. coli* membrane, Fig. 5A-C. A similar distribution was also observed for BcsZ and β-lactamase, as expected for soluble periplasmic proteins, Fig. 5A-C.

**Fig. 5.**
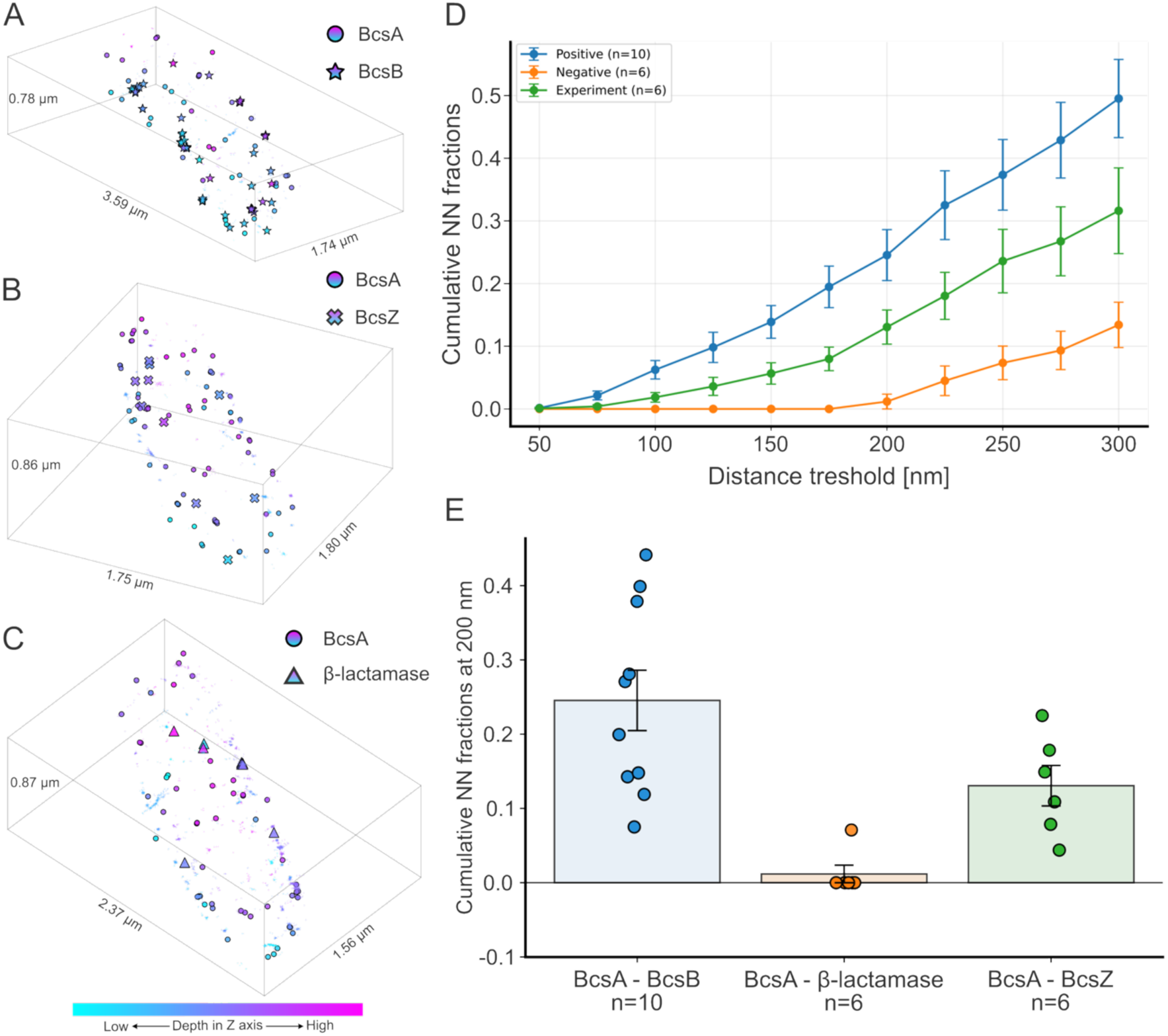
Cellular localization of BcsZ. (A-C) Representative MINFLUX centroids of localizations of BcsA together with BcsB, BcsZ, and β-lactamase colored based on the DCR assignment of fluorophores. (D) Nearest neighbor (NN) analysis of the localization centroids in the range of 0 – 300 nm for positive control (BcsB vs. BcsA), negative control (ß-lactamase vs. BcsA) and the experiment (BcsZ vs. BcsA). (E) Cumulative nearest neighbor fractions at a distance of 200 nm. All samples marked on the plot represent different technical replicates. Each group consists of samples originating from at least three different biological replicates.

A NN analysis of the co-localized fluorophores reveals the highest and lowest enrichment of BcsA neighbors for BcsB and β-lactamase, respectively, as expected for positive and negative controls, Fig. 5D, E and S10-S12. The BcsA-BcsZ pair, however, showed an intermediate distribution, with more nearest neighbor distances than the BcsA-β-lactamase but substantially less than the BcsA-BcsB controls. Because the measured distances include proteinaceous tags used for labeling and are calculated for centroids, they do not represent the exact distances between the farthest points of the interacting proteins. Nevertheless, the strong BcsA-BcsB enrichment and weak BcsA-β-lactamase association validate the analysis. The intermediate BcsA-BcsZ distribution indicates that BcsZ is unlikely to be a stable structural component of the Bcs complex in the same manner as BcsB is. However, transient interactions cannot be excluded.

## Discussion

Requiring cellulase activity for bacterial cellulose secretion is counterintuitive. However, it can be reconciled with a model by which mislocalized cellulose would stall cellulose biosynthesis, and its enzymatic degradation would allow biosynthesis to proceed. As mislocalized cellulose may be pEtN modified, BcsZ must be able to cleave both forms.

Indeed, BcsZ efficiently binds pEtN cellulose with preferred pEtN positioning at its −3 and +2 subsites. The moiety is excluded from binding to subsite −1 and −2, but may be tolerated at site +1, based on the glucose’s C6 hydroxyl position at this subsite. Crystallography indicates that additional glucosyl binding sites flanking the −3 to +2 positions can also accommodate pEtN modified units. This suggests that cellulose stretches with consecutive pEtN modifications can be recognized and cleaved, mainly due to limited protein interactions at the flanking −4 and +3 positions.

Polysaccharide processing during secretion is not limited to cellulose biosynthesis. Bacteria employ a spectrum of hydrolases, lyases, and epimerases as components of extracellular polysaccharide biosynthesis machineries. Similar roles in the degradation of mislocalized polymers have been proposed for the alginate lyase AlgL (29–31), the deacetylated-PNAG hydrolase PgaB (32, 33), and the bifunctional Pel deacetylase/hydrolase PelA (34, 35). However, the propensity of these enzymes to interact with other, especially outer membrane-associated, subunits of the biosynthetic machineries suggests additional, perhaps structural functions (34, 36). This has been proposed for the PslG hydrolase from *Pseudomonas aeruginosa* (37, 38).

Our MINFLUX localization of BcsZ and the cellulose biosynthetic machinery are consistent with a dynamic, perhaps random, localization of BcsZ in the periplasm. The fact that the cellulase shows increased proximity to BcsA compared to β-lactamase may be due to transient interactions with BcsC or the nascent cellulose polymer. Further, because the Bcs components are plasmid encoded and expressed from nonnative promoters, some co-localizations may be due to differences in expression levels.

Analyzing the cello-oligosaccharides released by BcsZ from *in vivo* produced pEtN cellulose identified unmodified, sparse, and densely modified regions. Consecutive pEtN modified glucosyl units demonstrate that both polymer sides are accessible to BcsG, the pEtN transferase. The presence of three BcsG subunits in the Bcs complex and the profound flexibility of its catalytic domain (16) likely ensure pEtN modification on both sides of the cellulose ribbon.

Cellulose biosynthesis is a rather slow process with estimated turnover rates in the millisecond regime (39). The in vitro catalytic rate of *E. coli* BcsG’s catalytic domain using synthetic pEtN donors and cello-oligosaccharides as acceptors has been reported to be even lower (k_cat_ ~3.8 x10^-6^ s^-1^). (40). This may explain the presence of multiple BcsG copies in the biosynthetic complex, as well as the stochastic occurrence of pEtN groups in the polymer, Fig. 6. Therefore, the rate of cellulose synthesis and translocation, accessibility of cellulose within the BcsB crown, and the conformational flexibility of BcsG’s catalytic domain may control the degree of cellulose modification.

**Fig. 6.**
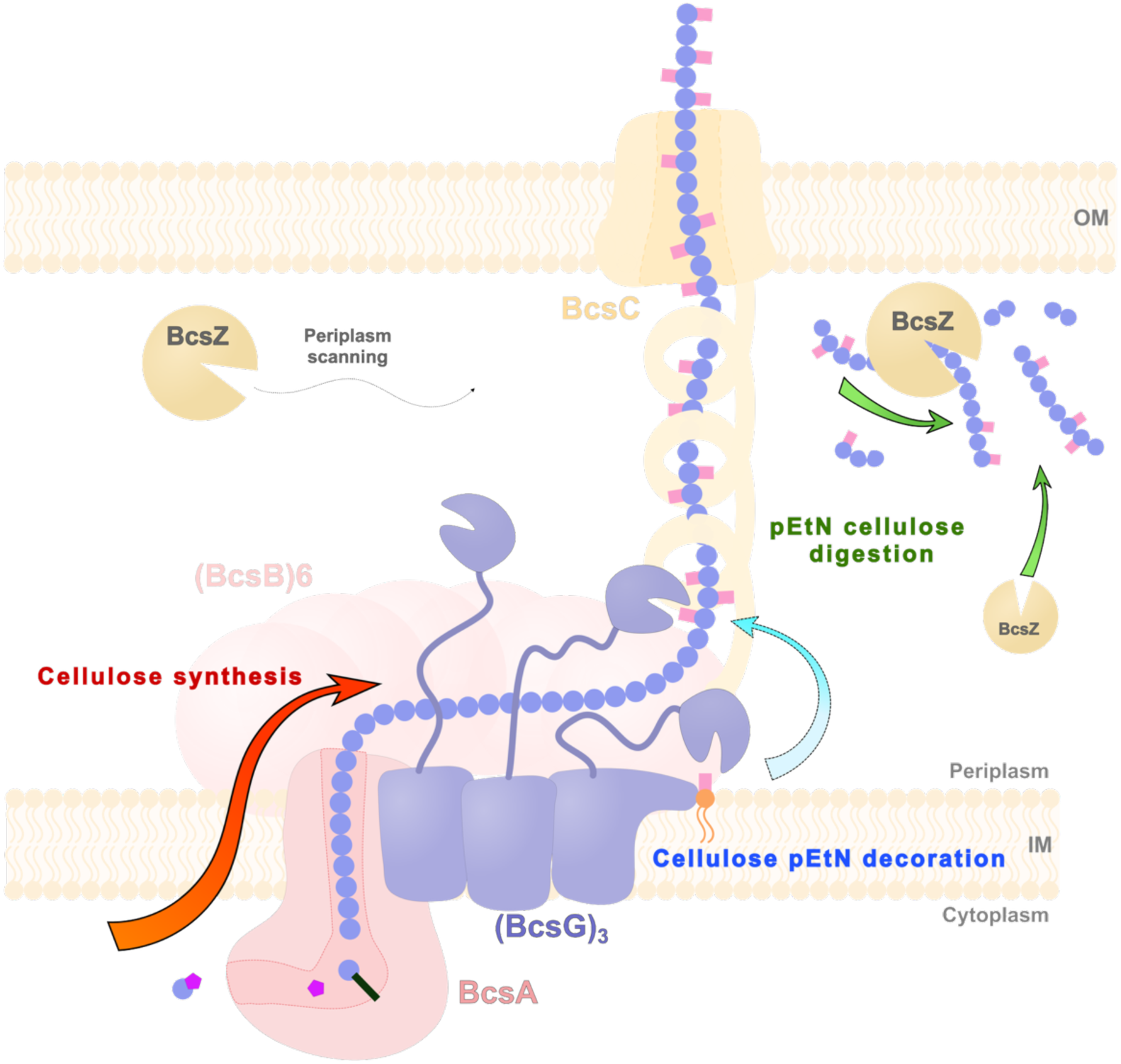
Cartoon representation of pEtN cellulose biosynthesis. The degree of pEtN modification is likely controlled by the cellulose biosynthesis rate, the accessibility of cellulose in the periplasm, as well as the catalytic rate of the pEtN transferase BcsG. BcsZ may transiently associate with the biosynthetic machinery to cleave mislocalized products accumulating in the periplasm.

Recent work analyzed the effects of pEtN cello-oligosaccharides on curli fibrillogenesis, based on the CsgA R5 peptide (23). Here, unmodified and pEtN-decorated cellohexaoses decreased the kinetics of R5 polymerization, thereby favoring the formation of fibrillar networks over random aggregation. Further, the pEtN distribution patterns of the compounds also influenced fiber growth rates and morphologies. This suggests that varying pEtN densities within cellulose polymers due to stochastic incorporation, different environmental conditions, or species traits could affect biofilm properties and virulence.

MINFLUX nanoscopy suggests that BcsZ does not stably interact with the Bcs macrocomplex. In our engineered *E. coli* system (15), all Bcs components are expressed from inducible vectors, likely resulting in higher than endogenous expression levels of the individual components. Because BcsZ fails to associate with BcsA even under those conditions, we conclude that its biological function is likely to patrol the periplasm and degrade mislocalized cellulose. A similar ‘housekeeping’ function may also be at play in other cellulose producing organisms (41).

## Materials and Methods

### Synthesis of phosphoethanolamine-modified cello-oligosaccharides

All synthetic oligosaccharides were prepared with automated glycan assembly following a reported procedure. For details, we refer to Ref. (22, 23).

### Protein expression and purification

Recombinant BcsZ-E55Q was expressed and purified as described previously (20). Briefly, *E. coli* C43(DE3) cells were transformed with a pET20b plasmid encoding the mature portion of BcsZ (aa 22–368) carrying the E55Q substitution, a C-terminal poly-His tag, and an N-terminal PelB signal sequence. Cells were cultured at 37 °C up to OD_600_ ≈ 0.4 and expression was induced with 0.5 mM isopropylthiogalactoside (IPTG) at 30 °C for 3 hours. The pelleted and microfluidized material was precleared by ultracentrifugation, and the soluble protein was purified by Ni-NTA affinity chromatography followed by size-exclusion chromatography (SEC) on a Superdex S200 column in 20 mM Tris-HCl, pH 7.5, and 100 mM NaCl. Following SEC, the purified protein was concentrated to 133 μM.

### Crystallization, ligand soaking, and data collection

Crystallization conditions for BcsZ-E55Q for soaking experiments were based on previously reported conditions for BcsZ (20). Crystals were grown at 20 °C using the sitting-drop vapor diffusion method. Crystallization drops consisted of 1 μL protein solution mixed with 1 μL reservoir solution containing 14–18% PEG3350, 100 mM sodium citrate buffer pH 7, and 40 mM NaCl. Well defined crystals appeared after ~4 days.

Ligand-bound complexes were obtained by soaking BcsZ-E55Q crystals with a series of synthetic phosphoethanolamine-modified cello-oligosaccharides, including APAAAP, APAAPA, AAPPAA, PAPAPA, APAAAA, PAAAAA and AAAAAP (22, 23). Soaking and cryoprotection were performed simultaneously using solutions containing 4.5 - 10 mM ligand, mother liquor supplemented with 2% higher PEG concentration relative to the crystallization condition, and stepwise increasing glycerol concentrations from 5–25%. For the thinnest crystals, 15% glycerol was sufficient for full cryoprotection. Cryoprotectant exchange was performed directly in the crystallization drop. 2 μL of soaking solution was added to the initial 2 μl crystallization drop containing the crystals. An additional 2 μL portion was added after 5 minutes of incubation, increasing the drop volume to 6 μL, after which 4 μL of solution was removed. This process was repeated sequentially while increasing glycerol concentration in 5% increments up to a final concentration of 15 - 25%. Following soaking and cryoprotection, crystals were flash cooled in liquid nitrogen.

BcsZ crystals bound to unmodified cellohexaose or -pentaose were obtained in the presence of 20-30 % PEG550 MME, 0.1 M Tris pH 8.5, in the presence of 2.5 mM cellohexaose or –pentaose. The crystals were cryoprotected and harvested as described above in the presence of 2.5 mM of the ligand.

Diffraction datasets were collected at the AMX beamline at the Brookhaven National Laboratory NSLS-II synchrotron (Brookhaven, NY, USA). Data were collected in both standard and vector mode using fine slicing with an oscillation range of 0.2° per image (0.1° for the complex with the APAAAP ligand). Exposure time varied between 0.005 to 0.01 s at a beam transmission of 10 - 25%. All data were collected on an EIGER detector.

### Data processing, structure solution, and refinement

Datasets were processed to resolutions ranging from 1.4–2.2 Å using XDS (42). Scaling, merging, and symmetry analysis were performed with AIMLESS (43) and POINTLESS (44) from the CCP4 package (45). Structures were solved by molecular replacement using Phaser-MR (46) with the previously deposited BcsZ structure (PDB ID: 3QXQ) as the search model. Model building and refinement were performed using COOT (47) in combination with both REFMAC5 (48) and Phenix (49). After refinement in the absence of a modeled ligand, well-resolved additional and non-proteinaceous densities allowed substrate modeling for most of the tested compounds. Structural visualizations were prepared in PyMOL (50). Software packages used in this project were curated by SBGrid (51).

### Comparative genomic analysis of BcsZ

BcsZ/GH8 homologs were obtained from an eggNOG-derived ortholog dataset (52), and one representative candidate per taxon was selected based on protein length and annotation, yielding 752 sequences. NCBI genome assemblies were identified using NCBI datasets, and candidate proteins were mapped to GenBank (53) CDS annotations using locus- and protein-specific identifiers. For successfully mapped targets, genomic neighborhoods comprising ±10 CDSs were extracted and screened for BcsA, BcsB, BcsC, and BcsG using a custom rule-based annotation pipeline with curated gene aliases and product descriptions. Medium- or strong-confidence assignments were used for classification. The primary dataset comprised 220 BcsZ sequences associated with at least two of the three cellulose-core components BcsA/B/C; an A+B+C subset of 152 sequences was analyzed independently as a higher-confidence control. Protein sequences were aligned with MAFFT v7.310 (54). The position equivalent to Arg177 of *E. coli* BcsZ (b3531) was identified from the alignment and extracted for each sequence.

### Electrospray ionization mass spectrometry

#### Testing MS-compatible buffers for BcsZ

To identify ammonium acetate concentrations compatible with BcsZ-WT activity and downstream MS analysis, cellopentaose hydrolysis in varying conditions was assessed by polysaccharide analysis using carbohydrate gel electrophoresis (PACE) (55). Reaction mixtures contained 20 μM cellopentaose in 100 μL ammonium acetate, pH 6.0, at concentrations ranging from 100 to 600 mM in 100 mM increments. A reference reaction used standard 20 mM Tris-HCl, pH 7.5, and 100 mM NaCl buffer. Reactions were initiated by adding 1.5 μL BcsZ-WT at 1.9 mg/mL, (enzyme final concentration of approximately 0.67 μM), and were incubated for 16 hours at 4 °C on a rocker. Untreated 20 μM cellopentaose in water was included as a negative control. Following incubation, the reaction mixtures, untreated substrate, and a cello-oligosaccharide standard mixture containing glucose through cellohexaose were dried by vacuum centrifugation. Samples were resuspended in 5 μL ANTS labelling solution containing equal volumes of 15% (v/v) acetic acid, DMSO, 0.2 M 2-picoline borane in DMSO, and 0.2 M 8-aminonaphthalene-1,3,6-trisulfonic acid disodium salt in 15% (v/v) acetic acid. Samples were derivatized overnight at 37 °C in the dark, dried by vacuum centrifugation, and resuspended in 2.5 μL 6 M urea. The entire sample volume was loaded onto 240 × 180 × 0.75 mm polyacrylamide gels containing a 10% stacking gel and a 20% resolving gel prepared with a 29:1 acrylamide-to-bis-acrylamide ratio in 0.1 M Tris-borate buffer, pH 8.2. Electrophoresis was performed in the same buffer at 200 V for 30 min, followed by 1,000 V for 2 h. NTS-labelled carbohydrates were visualized by in-gel fluorescence using excitation at 365 nm and detection at approximately 515 nm.

#### Native mass spectrometry (nMS) binding assay

Purified BcsZ-E55Q samples were buffer exchanged into 200 mM aqueous ammonium acetate (pH 6.0) using a 10 kDa cut-off Amicon 0.5 ml microconcentrator (EMD Millipore, Billerica, MA). Stock solutions of cellohexaose (C6) and phosphoethanolamine (pEtN) modified cellohexaose ligands (5 mM) were prepared by dissolving known amounts of the solid compounds in milliQ water. Both protein and ligand solutions were stored at −20 °C until used. The dissociation constant (*K*_D_) for each BcsZ-ligand interaction was measured using a direct nMS assay performed in a titration format and using a protein reference method for correction of nonspecific protein-ligand binding, as described elsewhere (56, 57). Aqueous ammonium acetate (200 mM, pH 6) solutions of the BcsZQ55N mutant (4 mM), reference protein lysozyme (1 mM), and individual ligands, at concentrations ranging from 10 to 300 mM, were prepared at room temperature and loaded into standard uncoated nanoflow electrospray ionization (nanoESI) borosilicate glass emitter (cat. No: ESIU-001, AB Glycomics, Edmonton, Canada). Binding measurements were performed in positive mode using a Q Exactive Orbitrap (Classic) mass spectrometer (Thermo Fisher Scientific, Bremen, Germany) equipped with a modified nanoESI source. A voltage of approximately 0.8 kV was applied to a platinum wire inserted inside the emitter and in contact with the sample solution. The inlet capillary of the MS was heated to 120 °C, the S-lens RF level was set at 100, automatic gain control target was set at 1 × 10^6^ with a maximum injection time of 100 ms. All MS data were acquired and processed using Thermo Xcalibur 4.2 software.

The *K*_D_ value for the 1:1 complex of each BcsZ-Q55N (P) and each ligand (L) was determined by fitting equation S1a to the concentration-dependent fraction (*R*/(*R*+1)) of the ligand-bound BcsZ protein (PL) (after correction for non-specific ligand binding using the reference protein method) (56):

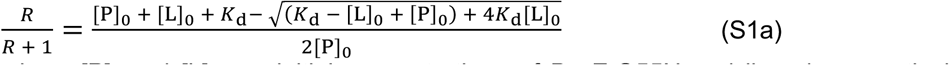

where [P]_0_ and [L]_0_ are initial concentrations of BcsZ-Q55N and ligand, respectively, and *R* is calculated form the corresponding total ion abundances (*Ab*) of ligand-bound to free BcsZ-E55Q protein (eq S1b):

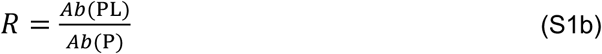

#### Analysis of enzyme kinetics using time-resolved nMS

Relative initial rates (*v*_0,rel_) for the BcsZ WT-catalyzed hydrolysis of the investigated substrates were determined using time-resolved nMS. Reactions were performed directly in a nanoESI emitter and continuously monitored by nMS, as described previously (58). Reaction mixtures containing BcsZ WT (0.15, 0.6, or 3 μM), standard substrate C6 (4 mM), substrate (4 μM), α-cyclodextrin (αCD, 1 μM), and 200 mM aqueous ammonium acetate (pH 6.0) were prepared in Eppendorf tubes, vortexed, and transferred to the emitter. Mass spectra were acquired continuously beginning at 3 min after reaction initiation, corresponding to the minimum time required for sample preparation and loading into the emitter. All measurements were performed at 22.5 °C. Three technical replicates were acquired and yielded similar results. Mass spectra were averaged over 1 min intervals and used to construct progress curves (normalized substrate concentration versus time). To account for potential changes in substrate response factors during the reaction, α-cyclodextrin (αCD), which is not hydrolyzed by BcsZ, was used as an internal reference. The abundances of the substrate and αCD ions were determined using the in-house SWARM software package (59). The fraction of intact (unreacted) substrate (*Fr*) was calculated using eq S2a, where *R*_0_ and *R*_t_ represent the abundance ratio of the substrate (*Ab*(Sub)) to αCD (*Ab*(αCD)) at the start of the reaction and at any time point (t), respectively (eqs S2b,c).

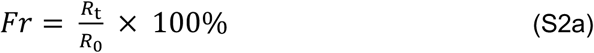

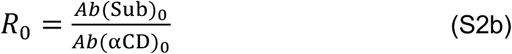

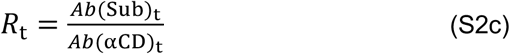

Initial rates of the reaction were obtained from the linear fit of the initial linear portion of the reaction progress curve (plot of the *Fr* versus time). Relative initial rates of the substates (n_0,rel_ were calculated using eq S3:

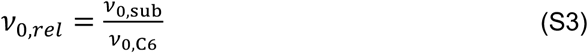

where *v*_0,sub_ and *v*_0,C6_ are initial rates of substrate and C6, respectively, that were measured simultaneously (substrate, C6, aCD, and enzyme were present in the same solution).

Time-resolved plots of enzymatic product distributions were generated from separate kinetic experiments containing only BcsZ WT and substrate. Product abundances were obtained directly from the corresponding time-resolved mass spectra.

#### nMS analysis of pEtN-modified cellulose fragments obtained by BcsZ WT hydrolysis of E. coli produced cellulose

*E. coli* C43(DE3) expressing the Bcs complex from inducible plasmids were grown as described previously (15), with modifications to culture scale and cell preparation. Two types of samples were used in the experiment, freshly produced pEtN cellulose still entangled with intact flocculent cells, and the same pEtN cellulose but purified according to the published protocol (15). An aliquot of each cell or purified cellulose sample was gently resuspended in 100 μL of 200 mM aqueous ammonium acetate (pH 6.0). BcsZ WT enzyme was then added, and the resulting mixture was incubated overnight at room temperature. Following centrifugation at 2000 × g, the supernatant was collected and analyzed by nMS. The compositions of the detected pEtN-modified cellulose species were assigned based on their measured molecular masses, as described previously (60). As a control, nMS analysis was also performed on cell samples that were not treated with BcsZ WT.

### MINFLUX nanoscopy

#### Expression and preparation of cells for imaging

Bcs components were expressed as described above with the following alterations to the original plasmids: His-tags were removed from all genes except BcsA and a Halo tag was fused to the C-terminus of BcsZ. All changes to vectors were made using the NEB Hi-Fi assembly methodology. *E. coli* L21 C43(DE3) cells were co-transformed by electroporation with the pETDuetEcBcs_A-12His_nSS-Strep-B_Adra, pACYCDuet_EcBcs_PelB-C_Z-HALO_F_G-FLAG, and pCDF_EcBcs_R_Q_HA-E plasmids and plated on LB agar supplemented with ampicillin, chloramphenicol, and streptomycin, as described before, (15)

Scrapes of colonies were used to inoculate 5 mL LB starter culture containing antibiotics and grown at 37 °C for 9 h with shaking. Expression cultures were prepared by inoculating 100 mL TB-M-80155 autoinduction medium with starter culture (initial OD = 0.07). The medium consisted of TB (24 g/L yeast extract and 12 g/L tryptone) supplemented with antibiotics, 20× M salts solution (1 M NH_4_Cl, 0.5 M Na_2_HPO_4_, 0.5 M KH_2_PO_4_, and 0.1 M Na_2_SO_4_), and 40× 80155 autoinduction solution containing glycerol, glucose, and lactose (15).Cultures were grown overnight at 30 °C with shaking at 220 rpm. After 12 h, the temperature was reduced to 25 °C and IPTG was added to a final concentration of 4 mM. Expression was continued for an additional 4 h before cells were harvested by centrifugation at 2,000 rcf for 20 min at 4 °C.

Following harvesting, cells were washed three times with PBS. For each wash, cell pellets were resuspended in 100 mL PBS distributed between two 50 mL centrifuge tubes and centrifuged at 1,500 rcf for 15 min at 4 °C. After the final wash, cells were resuspended in PBS to an OD_600_ of 10, aliquoted into 50 μL portions, flash-frozen in liquid nitrogen, and stored at −80 °C.

#### Congo red binding and fluorescence assay

To assess whether fusion of the HALO domain to BcsZ affects its function, Congo Red (CR) accumulation by macrocolonies expressing BcsZ-HALO was compared with that of cells expressing wild-type or catalytically inactive BcsZ. The assay was performed as described previously (16). Briefly, *E. coli* cells transformed with the three plasmids described above were cultured at 37 °C in LB medium with shaking, normalized to an OD_600_ of 1.0, spotted (5 μL) onto NaCl-free LB agar supplemented with 25 μg/mL CR, 250 μM IPTG, ampicillin, chloramphenicol, and streptomycin. Plates were incubated at room temperature for 48-56 hours, and CR fluorescence associated with colonies producing pEtN modified cellulose was visualized using a G:Box Chemi-XX6 imaging system (Syngene) at 497/614 nm excitation/emission. A CR accumulation phenotype comparable to that of BcsZ-WT was taken as evidence that the BcsZ-HALO fusion retained BcsZ function.

#### MINFLUX sample preparation and labeling

Glass coverslips were glow-discharged for 1 min 30 s at 25 mA. Plasma cleaned coverslips were then coated with 200 µL poly-L-lysine for 30 min in a humid chamber, followed by three brief washes with PBS. 4% solution of gold nanobeads in PBS was applied to the coverslips and incubated for another 30 min in a humidity chamber. Excess beads were removed by three PBS washes. Cell aliquots were thawed, diluted to an OD_600_ of approximately 1.0, and washed three times with PBS by brief centrifugation at 1,500 rcf. A 100 µL aliquot of the washed cell suspension was added to each coverslip and incubated for 30 min to allow cell attachment to the poly-L-lysine surface. Coverslips were then washed three times with PBS to remove unattached cells.

Cells were fixed with 4% paraformaldehyde (PFA) in PBS for 15 min at room temperature in a dark humidity chamber. After fixation, coverslips were rinsed three times with PBS. Residual PFA was quenched by incubation with 50 mM ammonium chloride in PBS for 30 min, followed by three additional PBS washes.

Cells were permeabilized with 0.03% Triton X-100 for 4.5 min. Coverslips were then washed five times with PBS for 30 min per wash on an orbital shaker in an aluminum foil-covered six-well cell culture dish. Samples were then blocked with BlockAid for 15 min at room temperature prior to labeling.

Fluorescent probes were diluted in PBS containing 20% BlockAid and applied to the samples for 2 h at room temperature in the dark. The probes used in this study included, anti-His Alexa Fluor 647 antibody (BcsA labeling), HaloTag Alexa Fluor 660 (BcsZ labeling), streptavidin-FLUX680 (BcsB labeling), and anti-ß-lactamase Alexa Fluor 680 antibody (ß-lactamase labeling). After labeling, coverslips were washed five times with PBS over 1 hour on a rocker while protected from light.

Anti-His Alexa Fluor 647 and Streptavidin FLUX 680 were purchased as ready-to-use products and both used in concentrations of 3-5 µg/mL. Labeling of BcsZ was done by charging the BcsZ-fused HALO tag with free haloalkane Alexa Fluor 660 at 200-300 nM concentration. Commercially available anti-ß-lactamase antibodies in concentration of 1 mg/mL were reacted with a 2-times molar excess of NHS Alexa Fluor 680 for 1 hour at 4 °C in the dark. The labeled sample was then dialyzed four times against PBS at 20,000-fold dilution factor using 50 kDa cutoff dialysis membranes.

Coverslips with labeled cells were mounted on dented glass slides in embedding medium of glucose oxidase/catalase (GLOX; 100 U/mL glucose oxidase, 1,200 U/mL catalase in 10 mM NaCl, 50 mM Tris pH 8, 2.5% glycerol, and 10% (w/v) d-glucose) buffer. The buffer also contained cysteamine (MEA) at varying concentrations from 14 to 18 mM. To assure proper oxygen depletion, samples were sealed with silicone glue.

#### MINFLUX data acquisition and processing

MINFLUX measurements were performed on a commercial MINFLUX microscope (Abberior Instruments GmbH), based on the setup described by Schmidt et al. (61). The instrument was equipped with a 100×/1.4 NA oil-immersion objective, a 640 nm continuous-wave excitation laser for imaging Alexa Fluor 647, Alexa Fluor 660, Alexa Fluor 680, and FLUX680 labeled targets in confocal and MINFLUX modes, a 488 nm pulsed laser for confocal pre-imaging, and a 405 nm continuous-wave laser for fluorophore activation.

Two-color MINFLUX imaging was performed using an adapted 3D imaging sequence provided by Abberior Instruments. A 640 nm excitation power of 3 to 5 µW was used during the first localization iteration and was gradually increased in subsequent iterations to 40 µW in the final iteration. Single-molecule fluorescence emission was collected between 650 and 720 nm with the pinhole set to 0.83 Airy Unit (AU). The 405 nm activation power was adjusted throughout each acquisition within the nW range to maintain a sparse and approximately constant single-molecule detection rate. For each sample, confocal images were acquired before MINFLUX imaging using a 488 nm excitation power of 2 µW, a pixel size of 50 nm, a dwell time of 10 µs, and an emission detection range of 500–550 nm. Sample drift was actively corrected using back-scattered light from gold beads illuminated with a 980 nm widefield laser. The microscope was operated using Imspector software (Abberior Instruments GmbH, v16.3) with MINFLUX acquisition drivers.

MINFLUX localizations were processed using custom Python scripts employing pandas, NumPy, and SciPy libraries. Localizations were first filtered using the 9^th^ iteration EFO (effective frequence at offset) range of 0-100,000 and grouped by trace identifier (tid). Only traces containing more than three localizations were retained, and each trace was represented by its 3D coordinate-wise median centroid. Fluorophore populations were assigned using a common 2D (iterations 7 and 9) DCR (detector channel ratio) gates, low: with values in a range of 0.10–0.40 (Alexa Fluor 660, Alexa Fluor 680, and FLUX680), and high: with DCR values of 0.55–0.85 (Alexa Fluor 647).

Bidirectional nearest neighbor (NN) distances between the two centroid populations were calculated in three dimensions using the Euclidean method implemented in ce.cdist (62). For each centroid in population A, the distance to its closest centroid in population B was determined, and vice versa; the two sets of nearest-neighbor distances were pooled for subsequent analysis. Cumulative NN fractions were calculated as the proportion of bidirectional NN distances within 50 to 300 nm in 25-nm increments. The observed distributions were additionally compared to label-shuffle and toroidal-shift null models generated within individual imaging regions.

Fluorophore localizations were performed on at least five different cells from three biological replicates.

### AI use statement

Custom Python scripts for MINFLUX data processing and visualization were developed with assistance from ChatGPT (OpenAI; GPT-5.5 Thinking, accessed June 2026). The tool was used to assist code drafting and debugging. All analytical decisions, final code, parameter settings, and resulting outputs were reviewed by the authors and subjected to manual sanity checks, including visual inspection of localization classifications and centroids, comparison with positive and negative controls.

## Data availability

Coordinates and structure factors have been deposited at the Protein data Bank under accessing codes 37VP-W.

## Acknowledgments

We are grateful to Jessica Matthias from Abberior for advice on MINFLUX data collection and processing, as well as Coleman Weaver for assistance in crystallizing wild type BscZ. We also thank the staff of the Brookhaven National Laboratory AMX-17 beamline for assistance in diffraction data collection. We also thank_ Arunabh Athreya, Ruoya Ho, and Louis Wilson for productive scientific discussion and help with the project.

This research used resources 17-ID-1 (AMX) or 17-ID-2 (FMX) of the National Synchrotron Light Source II; a U.S. Department of Energy (DOE) Office of Science User Facility operated for the DOE Office of Science by Brookhaven National Laboratory under Contract No. DE-SC0012704. The Center for BioMolecular Structure is primarily supported by the NIH, National Institute of General Medical Sciences through a Center Core P30 Grant (P30GM133893), and by the DOE Office of Biological and Environmental Research (KP1607011). J.R. and J.Z. were supported by NIH grant R35GM144130 awarded to J.Z. who also acknowledges the Howard Hughes Medical Institute, of which he is an investigator, for support.

J.H., T.T-E and M.D. thank the Max Planck Society, the Max Planck Queensland Centre on the Materials Science of Extracellular Matrices, and the German Federal Ministry of Education and Research (BMBF, grant number 13XP5114), for financial support.

J.S.K. acknowledges the Natural Sciences and Engineering Research Council of Canada for support.

This article is subject to HHMI’s Immediate Access to Research policy, which requires that this article be made publicly available as initial and revised preprints deposited on a designated preprint server under a CC BY 4.0 license.

## Author Contributions

J.R and J.Z. conceptualized the project. J.R. performed all crystallographic analyses and built and refined the BcsZ models. J.R. also performed and analyzed the MINFLUX experiments. J.H., T.T-E and M.D. synthesized the pEtN cellohexaose ligands. E.K., L.H. and J.S.K. performed the MS experiments and analyzed the data. J.Z. generated the first draft, and all authors edited the manuscript.

