## Supplemental figures S1-S12, Supplemental tables 1-2 for "Site-specific processing of phosphoethanolamine cellulose by the BcsZ cellulase reveals stochastic biofilm cellulose modification"

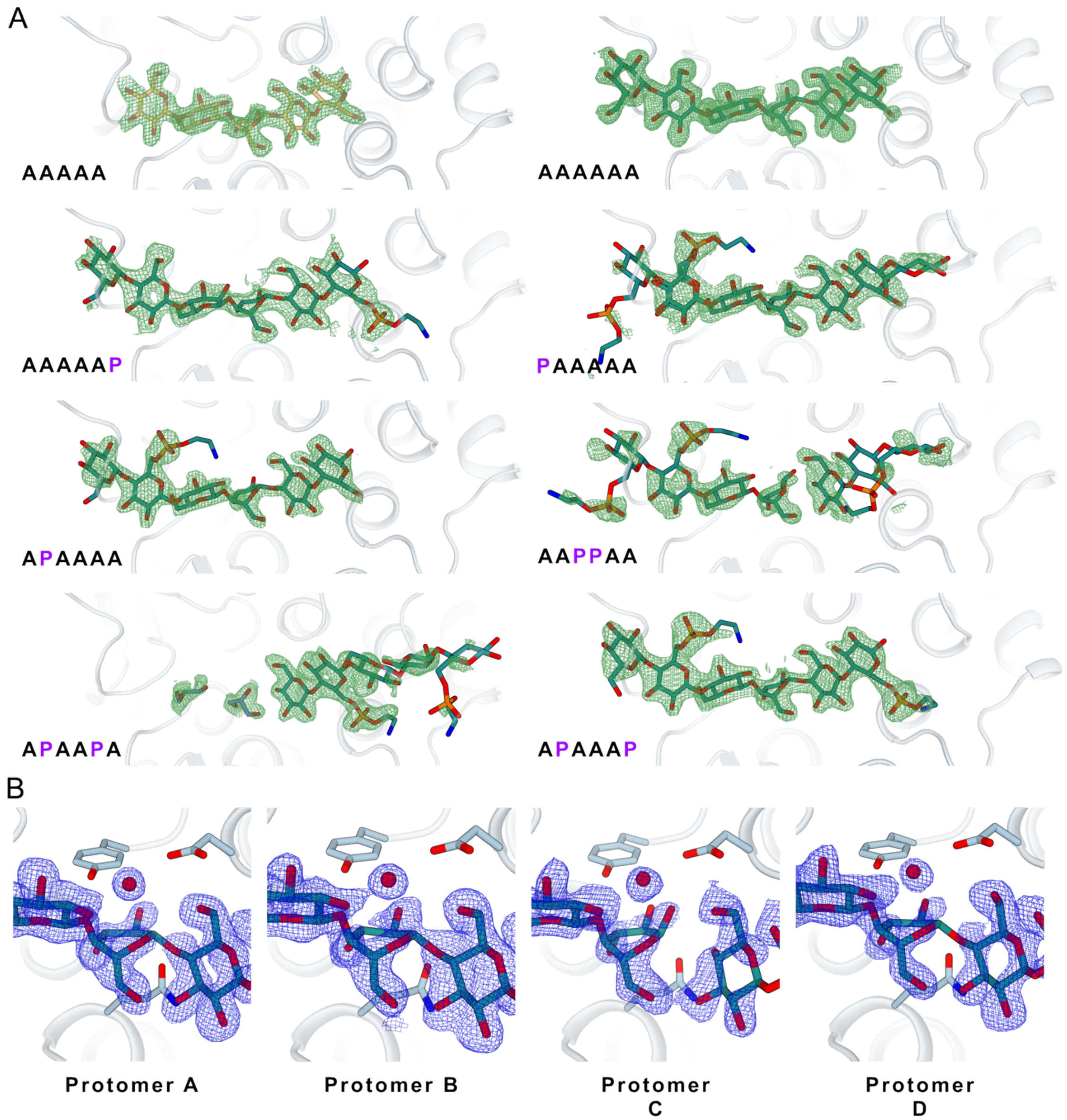


**Fig. S1.** Ligand electron density. (A) Unbiased FoFc difference density of the crystallographically resolved ligands shown as a green mesh and contoured at 3$\sigma$. (B) 2FoFc densities of the unmodified cellohexaose ligand bound to the four protomers of the crystallographic asymmetric unit, contoured at 1$\sigma$. The nucleophilic water is shown as a red sphere. Maps in protomers C and D show discontinuity of electron density in the areas of the scissile bond.

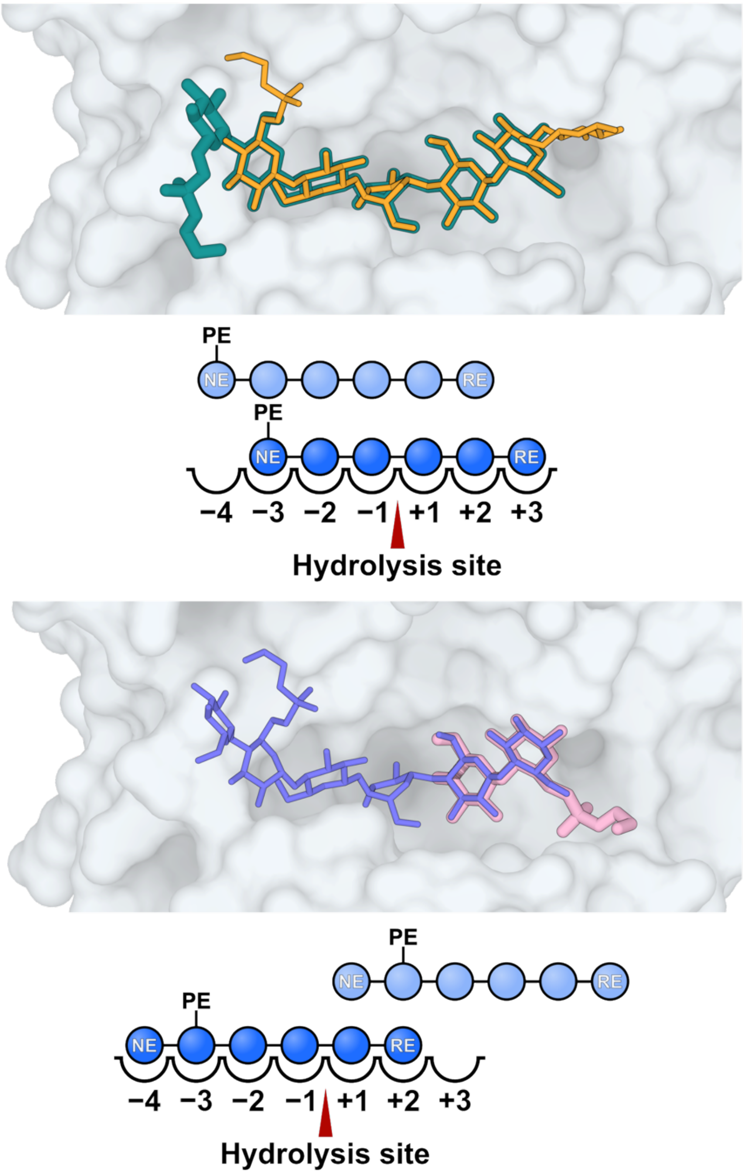


**Fig. S2.** Overlapping binding poses of the indicated ligands.

```
                                                                                          387
                                                                   350         .          .          .      V  .
400
  1 b3531|label=Escherichia_coli|BcsG=True                         VV----TVPGLGSMLLPG-KVGFA--ED---N---SWRFNPSYLPPTLAQY
  2 AXXA_09988|label=Achromobacter_insuavis|BcsG=False             VV----PLPGLGSMLLPA-PEGFAQ-PD-------LWRLNPSYLPVPVLRR
  3 AMBLS11_15740|label=Alteromonas_macleodii|BcsG=False           LV-----AHNKSVLLLPG-EYGFTS-EQ-AGKT--SLQLNLSYWIFPAITD
  4 GTGU_03217|label=Trabulsiella_guamensis|BcsG=True              VV----TVPGLGSMLLPG-KVGFV--DD---K---SWRFNPSYFPPQLTSY
  5 GKAS_01995|label=Kluyvera_ascorbata|BcsG=False                 VV----TVPGLGLMLLPG-RVGFA--EE---A---RWRFNPSYFPPQVAQY
  6 GBAG_4078|label=Buttiauxella_agrestis|BcsG=False               VI-----RFAGYQVMLPG-AKGFNL-NS-------YVNLNPSYFIFPAWQD
  7 EAE_05915|label=Enterobacter_aerogenes|BcsG=True               VA----DVPGLGPSVLPG-KVGFA--DD---K---GWRFNPSYLPPQLATY
  8 RG540_PA05620|label=Neorhizobium_galegae|BcsG=False            VI-----SAGGRTILLPG-AIGFRA-AD-RQD---GPVVNPSYWIFEALPV
  9 RG1141_PA10820|label=Neorhizobium_galegae|BcsG=False           VI-----SAGGRTILLPG-AIGFRA-ED-RQD---GPVVNPSYWIFEALPV
 10 RLEG12_17080|label=Rhizobium_leguminosarum|BcsG=False          VG-----SSQGRTLLMPG-TEGFTG-SD-RDD---GPVVNPSYWIYEAIPV
 11 HK44_004970|label=Pseudomonas_fluorescens|BcsG=True            LR----KLPGLGLMLLPG-DAGFD--NA---E---GWRLNASYLPPQLLAR
 12 Ser39006_00361|label=Serratia_sp.|BcsG=False                   II-----RFAGRTVMLPG-AVGFNK-NS-------YVILNPSYFLFPAWRD
 13 BN137_3848|label=Cronobacter_condimenti|BcsG=True              VV----NVPSLGDMLLPG-KVGFA--EP---T---LWRFNPSYLPPQLARY
 14 LH22_01685|label=Pantoea_rwandensis|BcsG=True                  VI-----TFGGHTVMLPG-AQGFNK-NS-------YVVLNPSYFLFAAWQA
 15 JIBP01000017_gene645|label=Lonsdalea_quercina|BcsG=True        TA----EIPDLGPVLLPG-KHGFA--SQ---T---SWTVNPSYLPPPLLVS
 16 KKY_993|label=Pelagibacterium_halotolerans|BcsG=False          LT-----EANGLTVILPG-AEGFSA-AA-RED---GPVVNPSYWVFEAFAV
 17 ATCR1_14111|label=Agrobacterium_tumefaciens|BcsG=False         VV-----QRGGRVLLLPA-ASGFDE-GD-RED---GPVVNPSYLIFEAFPV
 18 CM001373_gene1693|label=Fluoribacter_dumoffii|BcsG=True       VI----EIPGLGLMLLPG-KTGFN--FK---K---SWRLNPSYLPPQLFAR
 19 L284_04850|label=Novosphingobium_lindaniclasticum|BcsG=False   VI-----ERHGMSLLLPG-IMGFVN-GQ-------EVTLNPSYFAWPALDA
 20 BYI23_B008380|label=Burkholderia_sp.|BcsG=False                VV----DLPGLGPMLLPG-PHGFRT-GD-------VTRLNPSYLPLPLLRG
 21 TMO_1082|label=Tistrella_mobilis|BcsG=False                    IV-----EHAGRLMVLPG-QEGFR-----RDP---SVVVNPAYWVLPAFPV
 22 CIFAM_12_01020|label=Citrobacter_farmeri|BcsG=True             VV----TVPGLGSMLLPG-RIGFA--DA---T---SWRFNPSYLPPQLAQY
 23 EH105704_02_02780|label=Escherichia_hermannii|BcsG=False       VM-----NYAGYKVMLPG-AKGFDL-NS-------NINLNPSYFVFPAWRD
 24 EV102420_22_00730|label=Escherichia_vulneris|BcsG=True         VV----SVPGLGSMLLPG-KIGFA--EK---E---SWRFNPSYLPPQVAQY
 25 Salmuc_03686|label=Salipiger_mucosus|BcsG=False                IF----TASDGEPLLLPA-ARGFTT-DA-------GAIINPCYAMPLALTE
 26 NX02_16135|label=Sphingomonas_sanxanigenens|BcsG=False         IR-----QRGRRTLLVPG-LQGFLH-PD-------RTTINLSYYIWPALDA
 27 KB902315_gene3411|label=Wenxinia_marina|BcsG=False             IQPH--PDGSGRLVFTPA-AGGFQR-DE-------GLIVNPSYMMPLAMRE
 28 RCCGE510_00865|label=Rhizobium_sp.|BcsG=False                  VA-----SSDGRILLLPG-TEGFTG-SD-RED---GPVVNPSYWIYEAIPV
 29 OOA_15240|label=Providencia_burhodogranariea|BcsG=True         TM----DIPGLGWMLLPG-KVGFV--HG---N---EWKLNPSYQPPQLMAR
 30 PMI02_05305|label=Novosphingobium_sp.|BcsG=False               VV-----SRFGRQLLLPG-IVGFDE-PG-------QITLNPSYFVWPALDH
 31 PMI04_00616|label=Sphingobium_sp.|BcsG=False                   VL-----ERHGRHLLLPG-IQGFVT-AE-------AVTLNPSYYVWPALDA
 32 PMI09_00599|label=Rhizobium_sp.|BcsG=False                     VV-----SSGGRTLLLPG-ASGFDS-PD-RKD---GPIINPSYWVYEAIPI
 33 PMI40_00095|label=Herbaspirillum_sp.|BcsG=False                VK----VMPKLGPMLLPG-EFGFDA-GK---E---RWRLNPSYLPLPLLRR
 34 Q7S_00500|label=Rahnella_aquatilis|BcsG=False                  VI-----DYAGYKVMLPG-VKGFNQ-TS-------NVTVNPSYFIFPAWQA
 35 H650_14040|label=Enterobacter_sp.|BcsG=True                    VV----TVPGLGSMLLPG-KIGFA--EP---T---TWRFNPSYLPPQLASY
 36 UWK_01358|label=Desulfocapsa_sulfexigens|BcsG=False            LL-----EKFGKTLLLPG-YYGFAH-TD-------TVTVNPSYLVLPAFKV
 37 GWL_06950|label=Herbaspirillum_sp.|BcsG=False                  VV----DLPGLGKMVLPG-PQGFNQ-HN---Q---VWTLNPSYLPLPLLRR
 38 USDA257_c11960|label=Sinorhizobium_fredii|BcsG=False           VI-----DHGGRTLLLPG-VAGFSA-SD-RDD---GPVINPSYWVFEALPV
 39 LIG30_3521|label=Burkholderia_sp.|BcsG=True                    TA----VVPGLGLTLLPG-PVGFRP-AA---N---LWRVNPSYSPPQLIRG
 40 A464_3696|label=Salmonella_bongori|BcsG=True                   VV----NVPGLGSMLLPG-KIGFA--EA---N---SWRFNPSYLPPQLAQY
 41 AciX9_2051|label=Granulicella_tundricola|BcsG=False            VV----VVPELGTVLAPG-PKGFHP-DD---Q---TWLVNPSYMPPPVLVG
 42 HMPREF1476_00755|label=Sutterella_wadsworthensis|BcsG=True     VR----TVDGLGDVILPG-RVGFE--KN---G---LVKLNPSYYPLFILKR
 43 B195_17434|label=Pseudomonas_sp.|BcsG=True                     LR----TLPGLGLMLLPG-DVGFD--SA---Q---GWRLNPSYMPPQLLAR
 44 PP4_32190|label=Pseudomonas_putida|BcsG=True                   MR----KLPGLGVMLLPG-DYGFE--DA---R---GTRLNPSYLPLQLFDR
 45 BN77_1378|label=Rhizobium_mesoamericanum|BcsG=False            VI-----SSGGRTLLMPG-ASGFGS-PD-RKD---GPIVNPSYWIYEAIPI
 46 PA6_005_01360|label=Pseudomonas_alcaligenes|BcsG=False         TT----ELPGLGPVLLPG-PIGFSH-IG---H---LWRINPSYQPLPLLRR
 47 BAUC01000010_gene2349|label=Edwardsiella_hoshinae|BcsG=True    VV----ALPGFGMMLLPG-RHGFR--RG---A---LWTVNPSYLPPQILAR
 48 BV98_002173|label=Sphingobium_herbicidovorans|BcsG=False       VI-----ERFGRRLLLPG-LDGFSS-PN-------VVTVNPSYFIWPAFDQ
 49 N646_2958|label=Vibrio_alginolyticus|BcsG=True                 TV----NIAGVGTVLLPA-PTGFDA-DG-------QYRVNPSYVPLQLIAR
 50 DT73_09180|label=Mangrovibacter_sp.|BcsG=True                  VA----WLPGFGFLLLPG-NDGFT--SD---N---RWRLNPSYLPPQVIQR
 51 BUPH_05760|label=Burkholderia_phenoliruptrix|BcsG=False        TT----SLPGVGPMLLPG-PQGFRD-GG-------VTRLNPSYLPLPVLRA
 52 VIBNISOn1_450154|label=Vibrio_nigripulchritudo|BcsG=False      LK-----DTRGEFLLLPG-GYGFEF-SD-------KTIANLAYWVFPAFDD
 53 JPQP01000002_gene1045|label=Bordetella_trematum|BcsG=False     IT----ALPGFGAMLLPG-PAGFVQ-PG---P---LWRLNPSYLPVPLLRR
 54 AVS7_02037|label=Acidovorax_sp.|BcsG=True                      TA----DLPGLGPTLLPA-PRGFAP-EG---G---PWRLNPSYLPLQLLHG
 55 HY2_04810|label=Hyphomonas_sp.|BcsG=False                      VV-----ERQDRMILLPG-LNGFDH-DD-------LTIFNPAYFIWPAFRD
 56 HY29_15850|label=Hyphomonas_sp.|BcsG=False                     IV-----ERHDRMVLLPG-LNGFDH-DD-------RTVFNPAYFIWSAFQD
 57 H009_21641|label=Agrobacterium_tumefaciens|BcsG=False          VV-----QRGGRTLLLPA-ASGFDE-DD-RED---GPVVNPSYLIFEAFPV
 58 C266_21339|label=Pandoraea_sp.|BcsG=True                       TD----FVPGLGRTLLPA-PVGFHP-AK---D---LWRLNPSYVPMQVVRR
 59 RORB6_19920|label=Raoultella_ornithinolytica|BcsG=True         VT----DVPGFGPMVLPG-KVGFV--DD---K---GWRFNPSYLPPQLASY
 60 X805_06480|label=Sphaerotilus_natans|BcsG=False                VV----DVPGLGRMLLPG-PQGFVSADR---Q---QWRFNPSYSAVHQLRA
 61 H845_449|label=Gluconacetobacter_xylinus|BcsG=False            TM-----KVGSYEVLLPG-ATGFVT-KD-------AVTLNLSYYVMPSLMQ
 62 PKB_3048|label=Pseudomonas_knackmussii|BcsG=False              IK----DVPGLGRMLLPG-PVGYEH-PD---Q---LWRFNASYLPIPLLRR
 63 J139_13460|label=Pseudoalteromonas_agarivorans|BcsG=True      TA----YISGLGLTLLPA-PAGFEF-EN---N---TYKLNPSYSPLFIYQQ
 64 L581_1308|label=Serratia_fonticola|BcsG=True                   VA----NIPGLGLMLLPG-KMGFV--AE---D---RWRLNPSYLPPQLLAR
 65 IJ00_14200|label=Calothrix_sp.|BcsG=False                      TA----PGKEGKHYLLPGPKEAFLP-NP---N---TLYLNPSYLAPYAFRI
 66 OM33_08120|label=Pseudoalteromonas_sp.|BcsG=True               TE----FLPGLGLTVLPA-PYGFKV-DD---S---TWRLNPSYLPPFIFQR
 67 N018_21240|label=Pseudomonas_syringae|BcsG=False               IV----NLPGLGKMLLPG-PEGFAQ-PD---H---LWRLNPSYLPMPLLRR
 68 SRDD_06220|label=Serratia_sp.|BcsG=True                        VV----DIPDLGKMLLPG-KVGFV--AE---E---RWRLNPSYLPPQLLAR
 69 U771_01760|label=Pseudomonas_sp.|BcsG=False                    IV----RVPGLGKMLLPG-PVGYVH-GG---G---LWRFNPSYQVLAQLRR
 70 X548_01360|label=Stenotrophomonas_maltophilia|BcsG=True        IV----QIPTLGPMLLPG-PQGFI--ED---G---ATRVNPSYLPLPLLRR
 71 IE4771_CH01593|label=Rhizobium_etli|BcsG=False                 VG-----RSGGRTLLMPG-TAGFTG-SD-RED---GPVVNPSYWIYEAIPV
 72 GLUCORHAEAF1_08905|label=Gluconacetobacter_rhaeticus|BcsG=False TL-----QVGPYLVLLPG-GVGFAT-KD-------SVTLNLSYYVMPSLMQ
 73 X946_4479|label=Burkholderia_pseudomallei|BcsG=True            TA----TLPELGVALLPG-PEGFQP-AR---D---RWRLNPSYSPPQVIRG
 74 Z042_20780|label=Serratia_fonticola|BcsG=False                 LI-----KYAGYNVMLPG-VTGFRQ-AN-------SITINPSYFVFPAWEA
 75 OCH239_05500|label=Roseivivax_halodurans|BcsG=False            IA----ERSDGTPLLLPA-ADGFRT-AE-------GFIVNLAYSMPLALEE
 76 AT03_21190|label=Hafnia_alvei|BcsG=True                        VA----KLPGFGEMLLPG-RVGFA--HS---P---VWTVNPSYLPPQLMAR
 77 AZ34_04310|label=Hylemonella_gracilis|BcsG=True                VA----DIAPLGPTLLPG-PRGFAPGSGTEAG---SYRLNPSYVPPQLLRR
 78 KBK24_0100010|label=Burkholderia_sp.|BcsG=False                TT----TLPGLGPMLLPG-PQGFAN-GG-------VTRLNPSYLPLPVLRA
 79 PSNIH2_18010|label=Pantoea_sp.|BcsG=True                       VA----DLKGFGLMLMPG-KYGFI--EG---D---SWMINPSYLPLQLLAG
 80 PSNIH1_06205|label=Pantoea_sp.|BcsG=False                      VI-----DYAGHKVMLPG-ANGFNK-SS-------YVVLNPSYFLFAAWQA
 81 KO116_4182|label=Halomonas_sp.|BcsG=False                      TY-----EVAGFQVLLPG-IEGFRR-NE-------GADLNLSYWFIPALQD
 82 HR45_11435|label=Shewanella_sp.|BcsG=False                     VH-----EYAGHLVLLPG-LNGFTG-NG-------YIDINLSYWVMPALLA
 83 IV04_02745|label=Serratia_sp.|BcsG=False                       LV-----KYAGYNVLLPG-VSGFKQ-SN-------SITLNPSYFAFPAWQA
 84 JL39_00280|label=Rhizobium_sp.|BcsG=False                      VV-----GVKGQTVLLPA-TAGFGA-GE-RED---APVINPSYWIFEALPV
 85 JP74_04330|label=Devosia_sp.|BcsG=False                        LV-----QINGLRVTLPG-VEGFAV-GH-RDD---APVVNPSYWIWETRPV
 86 NC00_19225|label=Xanthomonas_sp.|BcsG=False                    VV----ALPGLGPMLLPG-RSGFV--EP---G---RWTLNPSYLPIQLLRR
 87 EDL933_4786|label=Escherichia_coli|BcsG=True                   VV----TVPGLGSMLLPG-KVGFA--ED---N---SWRFNPSYLPPTLAQY
 88 ATW7_09978|label=Alteromonadales_bacterium|BcsG=True           TA----YITGLGLSLLPA-PAGFEF-DN---E---RYKLNPSYSPLFIYQQ
 89 PP_2637|label=Pseudomonas_putida|BcsG=True                     VR----KLPGLGVMLLPG-DYGFE--DA---Q---GTRLNPSYLPLQLLDR
 90 Atu3307|label=Agrobacterium_fabrum|BcsG=False                  VV-----QRGGRTLLLPA-ASGFGE-GD-RAD---GPVVNPSYLIFEAFPV
 91 XAC3516|label=Xanthomonas_axonopodis|BcsG=False                VA----TLPGLGPMLLPG-RTGFV--DN---G---RWTLNPSYLPIQVLRR
 92 Dda3937_01994|label=Dickeya_dadantii|BcsG=False                II-----QFAGRTVMLPG-AYGFNK-NS-------YVILNPSYFLFPAWRD
 93 c4343|label=Escherichia_coli|BcsG=True                         VV----TVPGLGSMLLPG-KVGFA--ED---N---SWRFNPSYLPPTLAQY
 94 SULAZ_1375|label=Sulfurihydrogenibium_azorense|BcsG=False      VI-VKDNKLKDNYLLLPA-TYGFSN-EK--YD---IV-IFPSYYITFILKE
 95 BCAL1390|label=Burkholderia_cenocepacia|BcsG=True              TA----TVPGLGLTLLPG-PTGFKL-AD---G---QWRVNPSYSPPQVIRA
 96 PFLU_0303|label=Pseudomonas_fluorescens|BcsG=False             IV----RVPGLGKMLLPG-PVGYVH-AG---G---LWRFNPSYQVLAQLRR
 97 RL1648|label=Rhizobium_leguminosarum|BcsG=False                VV-----HASGHTLLLPG-SEGFAA-TD-RED---GPVVNPSYWIYEAIPV
 98 ECA4373|label=Pectobacterium_atrosepticum|BcsG=True            VV----NIPGLGVMLLPG-KVGFA--EK---E---SWRVNPSYLPPQLLAR
 99 SBG_3212|label=Salmonella_bongori|BcsG=True                    VV----NVPGLGSMLLPG-KIGFA--EA---N---SWRFNPSYLPPQLAQY
100 16504992|label=Salmonella_enterica|BcsG=True                   VV----NVPGLGSMLLPG-KIGFA--EA---N---SWRFNPSYLPPQLAQY
101 PSPTO_1029|label=Pseudomonas_syringae|BcsG=False               VV----NLPGLGKMLLPG-PEGFVQ-PD---H---LWRLNSSYLPVPLLRR
102 aq_1401|label=Aquifex_aeolicus|BcsG=False                      VP-V--CNGRRDYLFIPA-KEGYIK-NN-------IVSLNVVYYVPFIFRK
103 ACP_0076|label=Acidobacterium_capsulatum|BcsG=False            VA----NLPGLGSMLLPG-PTGFHP-NA---A---TWLLNPSYMPLPVVEG
104 BMAA1589|label=Burkholderia_mallei|BcsG=True                   TA----TLPGLGLVLLPG-PMGFRP-AR---D---AWRLNPSYSPPQAIRG
```

```
105 CV_2676|label=Chromobacterium_violaceum|BcsG=True        TA----DLPGLGLTLLPG-KDGFLF-SD---G---GARLNPSYLPPQLLAR
106 ZMO1086|label=Zymomonas_mobilis|BcsG=False               VL-----SGGGETILLPG-LQGFTG-TD-------YVILNFSYYIWPALKA
107 Rmet_2251|label=Cupriavidus_metallidurans|BcsG=True      TA----VLPGLGRTLLPG-PTGFHT-SP---D---VWRLNPSYVPLQVMRR
108 Bxe_B2042|label=Burkholderia_xenovorans|BcsG=False       MT----TLPGIGPMLLPG-PQGFKN-GG-------VTRLNPSYLPLPVLRA
109 Bcep1808_1350|label=Burkholderia_vietnamiensis|BcsG=False TA----SVPGLGLTLLPG-PTGFKL-AN---G---QWRVNPSYSPPQVIRG
110 MexAM1_META1p1169|label=Methylobacterium_extorquens|BcsG=False IL----FKDPHGPVLLPA-VSGFSA-RE-RAD---GPLINLSYWVFPAFQR
111 SMDB11_4255|label=Serratia_marcescens|BcsG=True          VA----DIPGLGLMLLPG-KVGFV--AE---D---RWRLNPSYLPPQLLAR
112 NJ56_13370|label=Yersinia_ruckeri|BcsG=True              VV----NIPGLGFMLLPG-KVGFV--HG---N---SWRLNPSYQPPQLLAR
113 PBPRA1722|label=Photobacterium_profundum|BcsG=True       TR----EVEGIGTVLLPG-KKGFDN-GK---G---QLRLNPSYVPLFLIKN
114 Arad_9976|label=Agrobacterium_radiobacter|BcsG=False     VV-----QVGGRTVLLPG-AEGFGA-TD-RDD---GPVINPSYWVYEALPV
115 Pnuc_1167|label=Polynucleobacter_necessarius|BcsG=True   IR-----KTNFGTVILPG-AFGFEK-PE-------GLKLNLSYWVFPAITE
116 VF_A0882|label=Vibrio_fischeri|BcsG=True                 TI----KVSGLGTVLLPG-KVGFVL-GK---N---HVRLNPSYVPLQLLTR
117 FP2506_04531|label=Fulvimarina_pelagi|BcsG=False         II-----DDRQGKIILPG-VEGFDA-DS-QPD---GPVVNLSYWVFPALAE
118 XCV3641|label=Xanthomonas_campestris|BcsG=False          VA----TLPGLGPMLLPG-RTGFV--DN---G---RWTLNPSYLPIQVLRR
119 85676513|label=Escherichia_coli|BcsG=True                VV----TVPGLGSMLLPG-KVGFA--ED---N---SWRFNPSYLPPTLAQY
120 NY99_03715|label=Xanthomonas_axonopodis|BcsG=False       VA----TLPGLGPMLLPG-RTGFV--DN---G---RWTLNPSYLPIQVLRR
121 Avin_05060|label=Azotobacter_vinelandii|BcsG=False       IK-----EHAGLTVLLPG-LQGFEQ-NG-------QLILNPSYLVMPAIRA
122 Bamb_1259|label=Burkholderia_ambifaria|BcsG=True         TA----SVPGLGLTLLPG-PTGFKL-AN---G---QWRVNPSYSPPQVIRG
123 RHE_CH01544|label=Rhizobium_etli|BcsG=False              VV-----RSAGRTLLMPG-SEGFGA-AD-RDD---GPVVNPSYWIYEAMPV
124 LPU83_3340|label=Rhizobium_sp.|BcsG=False                VV-----SSGGHTLLLPG-VSGFDS-SD-RKD---GPIVNPSYWVYEAIPI
125 Acry_0570|label=Acidiphilium_cryptum|BcsG=False          SV----AINGFGRVLIPG-ASDFP--DT---P---PVIVDPSYTPLFLARG
126 yinte0001_36830|label=Yersinia_intermedia|BcsG=True      VV----NIPGLGEMLLPG-KIGFN--DN---D---RWRLNPSYLPPQLLAR
127 DJ58_442|label=Yersinia_frederiksenii|BcsG=True          VV----NIPGLGEMLLPG-KIGFS--DD---D---RWRLNPSYLPPQLLAR
128 CN09_22095|label=Agrobacterium_rhizogenes|BcsG=False     VV-----QVGGRTVLLPG-AEGFGA-TD-RDD---GPVINPSYWVYEALPV
129 BAV2628|label=Bordetella_avium|BcsG=False                LA----DLPDFGPMLLPA-PFGFVQ-PD---G---IWRLNPSYLPIPVLRR
130 ECP_3631|label=Escherichia_coli|BcsG=True                VV----TVPGLGSMLLPG-KVGFA--ED---N---SWRFNPSYLPPTLAQY
131 Smed_5210|label=Sinorhizobium_medicae|BcsG=False         VV-----EHGGLTLLLPG-AQGFSA-AD-RAD---GPVINPSYWVFEAFPV
132 XFF4834R_chr34110|label=Xanthomonas_fuscans|BcsG=False   TA----TLPGLGPMLLSG-RTGFV--DN---G---RWTLNPSYLPIQVLRR
133 NG99_05400|label=Erwinia_typographi|BcsG=True            VI-----TFNGHTVMLPG-SQGFNK-TS-------YVVLNPSYFLFPAWRD
134 Riv7116_5437|label=Rivularia_sp.|BcsG=False              TF----TDSSEKRYLLPGPSDAFVP-NS---S---TIYLNPSYLAPYAFRI
135 YE4072A|label=Yersinia_enterocolitica|BcsG=True          VV----NIPGLGEMLLPG-KIGFN--NE---E---RWRLNPSYLPPQLLAR
136 Bmul_1925|label=Burkholderia_multivorans|BcsG=True       TA----NVPGLGLTLLPG-PTGFAL-AR---D---RWRVNPSYSPPQVIRA
137 Rleg2_1206|label=Rhizobium_leguminosarum|BcsG=False      VG-----SSQGRTLLMPG-TEGFTG-SD-RDD---GPVVNPSYWIYEAIPV
138 Lcho_2071|label=Leptothrix_cholodnii|BcsG=True           VI----ELPGLGTTLLPG-PQGFRR-GE---R---GAKLNPSYLPPQLLRW
139 Bphyt_5838|label=Burkholderia_phytofirmans|BcsG=False    MT----TLPGVGPMLLPG-PQGFKS-GG-------VTRLNPSYLPLPVLRS
140 Ent638_3928|label=Enterobacter_sp.|BcsG=True             VI----KVPGLGLMLLPG-NVGFT--EE---K---AWRFNPSYLPPQLANY
141 AciPR4_1395|label=Terriglobus_saanensis|BcsG=False       VV----SVPGIGTMLLPG-PTGFHP-DP---T---NYVLNPSYLPPQALAR
142 Mext_1369|label=Methylobacterium_extorquens|BcsG=False   IL----FKDPHGPVLLPA-VSGFSA-RE-RAD---GPLINLSYWVFPAFQR
143 M446_0105|label=Methylobacterium_sp.|BcsG=False          VL----FRSEGAALLLPG-MAGFSA-ED-RAD---GPVINLSYWIFPALAR
144 Mrad2831_4797|label=Methylobacterium_radiotolerans|BcsG=False LL----PRAPQGPLILPA-VAGFSA-ED-RAD---GPVVNLSYWVFPALDR
145 Mpop_1314|label=Methylobacterium_populi|BcsG=False       IL----FKDPHGPLLLPA-VSGFSA-RE-RAD---GPLINLSYWVFPAFQR
146 SJA_C2-05480|label=Sphingobium_japonicum|BcsG=False      IV-----QRHGFTLLLPG-LVGFAK-AD-------RVIVNPSYYVWPAIDA
147 ETA_33880|label=Erwinia_tasmaniensis|BcsG=False          VI-----VFGGRTLMLPG-AEGFNK-TS-------YVVLNPSYFVFPAWRD
148 B21_03332|label=Escherichia_coli|BcsG=True               VV----TVPGLGSMLLPG-KVGFA--ED---N---SWRFNPSYLPPTLAQY
149 HMPREF0189_00178|label=Burkholderiales_bacterium|BcsG=True VR----DVPTVGKIILPG-RVGFE--EK---G---VITINPSYYPPHILRR
150 EcolC_0186|label=Escherichia_coli|BcsG=True              VV----TVPGLGSMLLPG-KVGFA--ED---N---SWRFNPSYLPPTLAQY
151 RHECIAT_CH0001612|label=Rhizobium_etli|BcsG=False        VA-----RSAGQMLLLPG-SEGFTA-AD-RKD---GPVVNPSYWIYEAIPV
152 ESCAB7627_4529|label=Escherichia_albertii|BcsG=False     VV----TVPGLGSMLLPG-KVGFV--ED---N---TWRFNPSYLPPTLAHY
153 Cseg_3476|label=Caulobacter_segnis|BcsG=False            LV-----KQGNRTLLVPG-VDGFVN-AQ-------GMIFNPSYAIMPALDA
154 GU3_14400|label=Oceanimonas_sp.|BcsG=True                LH----WLPGLGLTLLPG-KQGFE--KA---E---GWKLNPSYLPLPLLAR
155 PanABDRAFT_3253|label=Pantoea_sp.|BcsG=False             VT-----NFGGYTVLLPG-AQGFNK-TS-------YVVLNPSYFLFQAWRE
156 ECTPHS_00774|label=Ectothiorhodospira_sp.|BcsG=False     LV-----DWHGRTILLPG-VDGFVH-GE-DGDTPARLVVNLSYWVFPALDM
157 PMI3103|label=Proteus_mirabilis|BcsG=False               LV-----KTENYSVLLPG-INGFKT-PE-------EIIINPSYFIFPAWKD
158 KP22_19965|label=Pectobacterium_betavasculorum|BcsG=True VV----NIPELGVMLLPG-KEGFA--EK---E---SWRVNPSYLPPQLLAR
159 Zmob_0234|label=Zymomonas_mobilis|BcsG=False             VL-----SGGGETILLPG-LQGFTG-TD-------YVILNFSYYIWPALKA
160 WSK_3365|label=Novosphingobium_sp.|BcsG=False            VV-----SRYDRQLLLPG-MIGFAE-AG-------QVTLNPSYFVWPALDH
161 Dd1591_4079|label=Dickeya_zeae|BcsG=False                II-----QFAGRTVMLPG-AYGFNK-NS-------YVVLNPSYFLFPAWRD
162 PC1_0074|label=Pectobacterium_carotovorum|BcsG=True      VV----NIPGLGVMLLPG-KVGFA--EK---E---SWRLNPSYLPPQLLAR
163 Pecwa_0071|label=Pectobacterium_wasabiae|BcsG=True       VV----NIPGLGMMLLPG-KVGFT--EK---E---SWRLNPSYLPPQLLAR
164 Zymop_0203|label=Zymomonas_mobilis|BcsG=False            VL-----SGGGETILLPG-LQGFVG-TD-------HVTLNFSYYIWPALKA
165 Dd703_3891|label=Dickeya_dadantii|BcsG=False             VV-----QFAGRTVMLPG-ANGFNK-NS-------YVILNPSYFLFPAWQD
166 Msip34_0042|label=Methylovorus_glucosetrophus|BcsG=False VK-----PSKWGPVILPG-SAGFEK-PA-------GQIVNLSYWIFPALND
167 Dd586_4057|label=Dickeya_dadantii|BcsG=False             II-----QFAGRTVMLPG-AYGFNK-NS-------YVVLNPSYFLFPAWQD
168 Pat9b_0118|label=Pantoea_sp.|BcsG=True                   VI-----TYAGHTVMLPG-VQGFNK-TS-------FIVLNPSYFLFPAWRE
169 S7A_04585|label=Pantoea_sp.|BcsG=True                    VA-----NFGGHTVLLPG-AQGFNK-GS-------FVVLNPSYFLFQAWHE
170 Varpa_3210|label=Variovorax_paradoxus|BcsG=False         VV----ALPGLGRMLLPW-PASVA--SG---P---VWRLNPSYMPLQLLQY
171 LG71_02375|label=Pluralibacter_gergoviae|BcsG=True       VI-----TFAGYRLMLPG-AQGFNL-GD-------KVNLNPSYFIFPAWQA
172 bglu_2g21310|label=Burkholderia_glumae|BcsG=False        VV----DLAGLGPMLLPG-PQGFVN-GD-------VTRLNPSYLPLPVLRG
173 EBL_c01500|label=Shimwellia_blattae|BcsG=True            VV----RVPGLGMVLLPG-KQGFV--QP---G---SWRLNPSYLPPQVLAR
174 GLX_25040|label=Komagataeibacter_medellinensis|BcsG=False TM-----QVGPYKVLLPG-GVGFVN-KD-------SVTLNLSYYVMPSLMQ
175 EpC_35980|label=Erwinia_pyrifoliae|BcsG=False            VI-----AFGGRTVMLPG-AQGFNK-TS-------YVVLNPSYFVFPAWRD
176 EbC_43980|label=Erwinia_billingiae|BcsG=True             VI-----TFGGQTVMLPG-VQGFNK-TS-------YVVLNPSYFLFPAWRD
177 Acaty_c0516|label=Acidithiobacillus_caldus|BcsG=False    VV----HLNGWGPVLTPG-DGPYWR-VD---D---GFVVNPSYLPPFVLAG
178 SVI_0127|label=Shewanella_violacea|BcsG=True             TL----EVPGIGNVLLPG-PQGFDK-PD---G---SYRLNPSYVPLQLLSR
179 ROD_42761|label=Citrobacter_rodentium|BcsG=True          VV----TVPGLGSMLLPG-KIGFA--EE---N---SWRFNPSYLPPQLAQY
180 Snov_3172|label=Starkeya_novella|BcsG=False              LV-----TDGSRLVLLPG-VSGFSS-SD-RPD---GPILNLSYWIFETFPV
181 BC1002_3562|label=Burkholderia_sp.|BcsG=False            VA----NLSGLGPMLLPG-AQGFRN-GG-------VTRLNPSYLPLPLLRA
182 MD26_13740|label=Pseudomonas_sp.|BcsG=True               VR----KLPGLGVMLLPG-DYGFE--DA---N---GTRLNPSYLPLQVLDR
183 EAMY_3602|label=Erwinia_amylovora|BcsG=False             VI-----AFGGRTVMLPG-VQGFNK-TS-------YVVLNPSYFVFPAWRD
184 VMC_35740|label=Vibrio_alginolyticus|BcsG=True           TV----NVAGVGTVLLPA-PTGFDA-DG-------QYRVNPSYVPLQLIAR
185 AciX8_2185|label=Granulicella_mallensis|BcsG=False       VS----HVHSF-PILLPA-QTGFEH-DG---M---T-VMNPSYMPLFLLQS
186 PVLB_00980|label=Pseudomonas_sp.|BcsG=True               VR----KLPGLGVMLLPA-DYGFE--SD---K---GWRLNPSYLPPQLFDR
187 D782_0205|label=Enterobacteriaceae_bacterium|BcsG=True   VV----NVPGLGSMLLPG-KIGFA--EA---N---SWRFNPSYLPPQLASY
188 VIBRN418_00275|label=Vibrio_sp.|BcsG=True                TV----TQTNGEVLLLPA-PHGFET-KA-------GYRVNPSYVPLQLIAR
189 Entcl_0212|label=Enterobacter_lignolyticus|BcsG=True     VV----TIPKLGAVLLPG-KVGFA--DD---K---GWRLNPSYLPPQLARY
190 PANA_0132|label=Pantoea_ananatis|BcsG=False              VT-----NFGGYTVMMPG-AQGFNK-NS-------YVVLNPSYFLFVAWQE
191 Pvag_3358|label=Pantoea_vagans|BcsG=True                 VT-----SFGGHTVLLPG-AQGFNK-TS-------YVVLNPSYFLFQAWRE
192 ambt_00585|label=Alteromonas_sp.|BcsG=False              FI-----ETEAFLVLLPG-EFGFKS-DE-SNDE--GIQLNLSYWVFPAIDA
193 ECL_04936|label=Enterobacter_cloacae|BcsG=True           VV----KVPGLGLMLLPG-KVGFV--DE---N---AWRFNPSYLPPQLASY
194 Rahaq_0102|label=Rahnella_sp.|BcsG=False                 VI-----DYAGYKVMLPG-VKGFNQ-TS-------NVTVNPSYFIFPAWQA
195 HMPREF9464_00791|label=Sutterella_wadsworthensis|BcsG=True VR----TVDGLGDVILPG-RVGFE--KN---G---LVKLNPSYYPLFILKR
196 HMPREF9465_00130|label=Sutterella_wadsworthensis|BcsG=True VR----TVDNLGSVVLPG-RVGFE--DN---G---TVKLNPSYYPLFLFKR
197 Acife_1587|label=Acidithiobacillus_ferrivorans|BcsG=False VV----TVPGIGVMLLPG-KEGFHK-NP---D---IYIFNLSYLPLPVVDA
198 Psesu_2413|label=Pseudoxanthomonas_suwonensis|BcsG=False VT----ELPGFGTMLLPG-NHGYV--HA---G---SWTLNPCYLPLFAFRR
199 TRICHSKD4_2567|label=Roseibium_sp.|BcsG=False            IR-----SPKGDLALLPG-KHGFTA-DD-RPD---GPIVNLSYWVFPAFDA
200 GRAQ_04047|label=Rahnella_aquatilis|BcsG=False           VI-----DYAGYKVMLPG-VKGFNQ-TS-------NVTVNPSYFIFPAWQA
201 B3C1_02320|label=Gallaecimonas_xiamenensis|BcsG=False    VR-----PYAGKLVLLPG-LEGFDG-GD-------YLDLNLSYWVMPAFRD
202 HMPREF9439_01882|label=Parasutterella_excrementihominis|BcsG=True VR----DVPTVGKIILPG-RVGFE--EK---G---VITINPSYYPPHILRR
203 Fraau_2357|label=Frateuria_aurantia|BcsG=True            AA----RIPGLGPMLLPG-RLGFA--GT---G---WWKLNPSYLPIQVLRR
204 PputGB1_3195|label=Pseudomonas_putida|BcsG=True          VR----KLPGLGVMLLPG-DYGFE--DA---Q---GTRLNPSYLPLQLFDR
205 VIBC2010_04654|label=Vibrio_caribbeanicus|BcsG=False     IR-----IVDGDTILLPG-NNGFET-PE-------STIVNPAYWIFPAFKE
206 HFRIS_021066|label=Herbaspirillum_frisingense|BcsG=False VV----DLPGLGKMLLPG-PQGFNQ-RN---A---LWTLNPSYLPLPLLRR
207 OsccyDRAFT_0771|label=Oscillatoriales_cyanobacterium|BcsG=False TVADYRTDAQQRRYLLPGPATSF-Q-KK---T---IIQLNPSYFSPASFRL
208 HGR_03147|label=Hylemonella_gracilis|BcsG=False          VV----DISPLGPTLLPG-PWGFAPGSGTEPG---SYRLNPSYVPPQLLRR
209 GEAM_3552|label=Ewingella_americana|BcsG=True            VI-----DYAGYKVMLPG-VSGFNQ-TS-------SITLNPSYFLFPAWQA
210 GLAD_01796|label=Leclercia_adecarboxylata|BcsG=False     VI-----SFAGYRVMLPG-AKGFNL-NS-------YVNLNPSYFIFPAWED
211 CF149_02829|label=Pseudomonas_sp.|BcsG=True              LR----KMPGLGLMLLPG-DVGFD--SP---K---GWRLNPSYLPPQLLAR
```

```
212 XVE_0432|label=Xanthomonas_vesicatoria|BcsG=False                 VA----ALPGLGPMLLPG-RSGFV--ET---G---RWTLNPSYLPIQLLRR
213 Terro_1433|label=Terriglobus_roseus|BcsG=False                    VA----VSPSLGTVLLPG-PTGFKP-QP---N---TMILNPSYSPPQVLAR
214 LV28_03070|label=Pandoraea_pnomenusa|BcsG=True                    TD----FVPGLGRTLLPA-PVGFHP-AK---D---LWRLNPSYVPMQVVRR
215 PAJ_3292|label=Pantoea_ananatis|BcsG=False                        VT-----NFGGYTVMMPG-AQGFNK-NS-------YVVLNPSYFLFVAWQE
216 VISI1226_01950|label=Vibrio_sinaloensis|BcsG=True                 TA----MIEGIGTILLPA-PVGFHL-EQ-------GYRVNPSYVPLQLLAR
217 Y88_1849|label=Novosphingobium_nitrogenifigens|BcsG=False         VV-----ERYDRHLLLPG-RTGFFT-SE-------AITLNPAYFVWPALDA
218 RGCCGE502_00240|label=Rhizobium_grahamii|BcsG=False               VI-----NSGGRTLLLPG-ASGFDS-PD-RKD---GPIINPSYWVYEAIPV
219 Cal7507_3086|label=Calothrix_sp.|BcsG=False                       SA----PGYGGKRYLLPGPKAAFIP-NE---S---TLYLNPSYLAPYAFRI
220 bgla_2g28050|label=Burkholderia_gladioli|BcsG=False               VA----DLAGLGPMLLPG-PQGFVE-GD-------VTRLNPSYLPLPVLRA
```

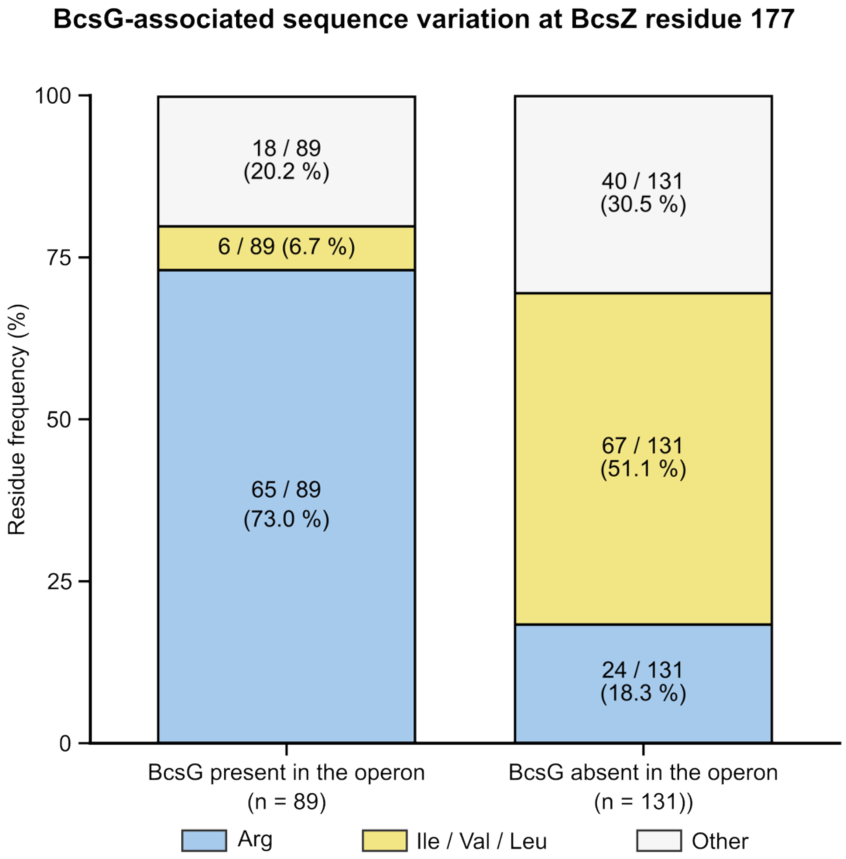


**Fig. S3.** Multiple sequence alignment of BcsZs from pEtN-producing bacteria generated in MAFFT. Bottom panel: Correlation of an R or I/V/L residue at position 177 (*E. coli* numbering) with the presence of a bcsG gene in the bcs operon. The presence of a bcsG gene is assumed to indicate the production of pEtN cellulose by the Bcs system.

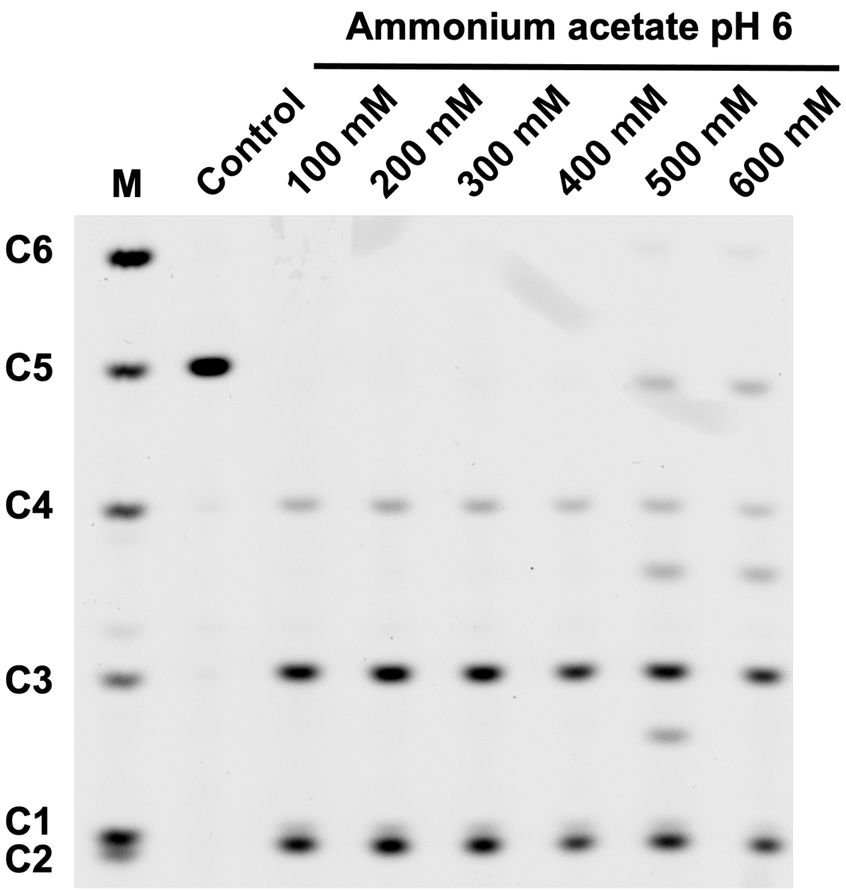


**Fig. S4.** Analysis of BcsZ's catalytic activity on the presence pf ammonium acetate. BcsZ's catalytic activity was analyzed in the presence of the indicated concentrations of ammonium acetate by polysaccharide analysis by carbohydrate electrophoresis (PACE) M: carbohydrate marker including mono to hexa-glucosides (C1-C6).

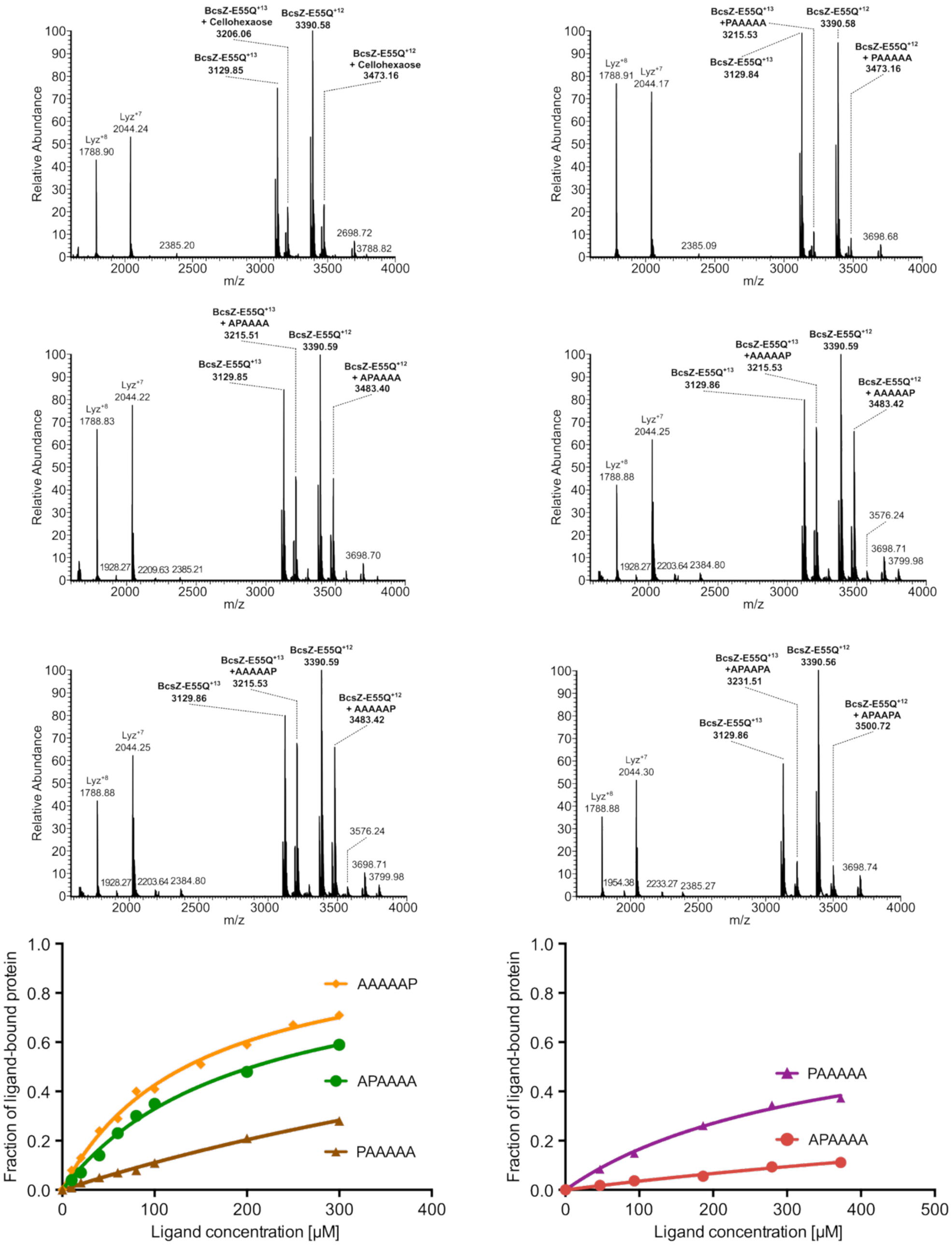


**Fig. S5.** Representative mass spectra for the indicated BcsZ-ligand complexes. Bottom panels: Quantification of ligand bound species depending on ligand concentration.

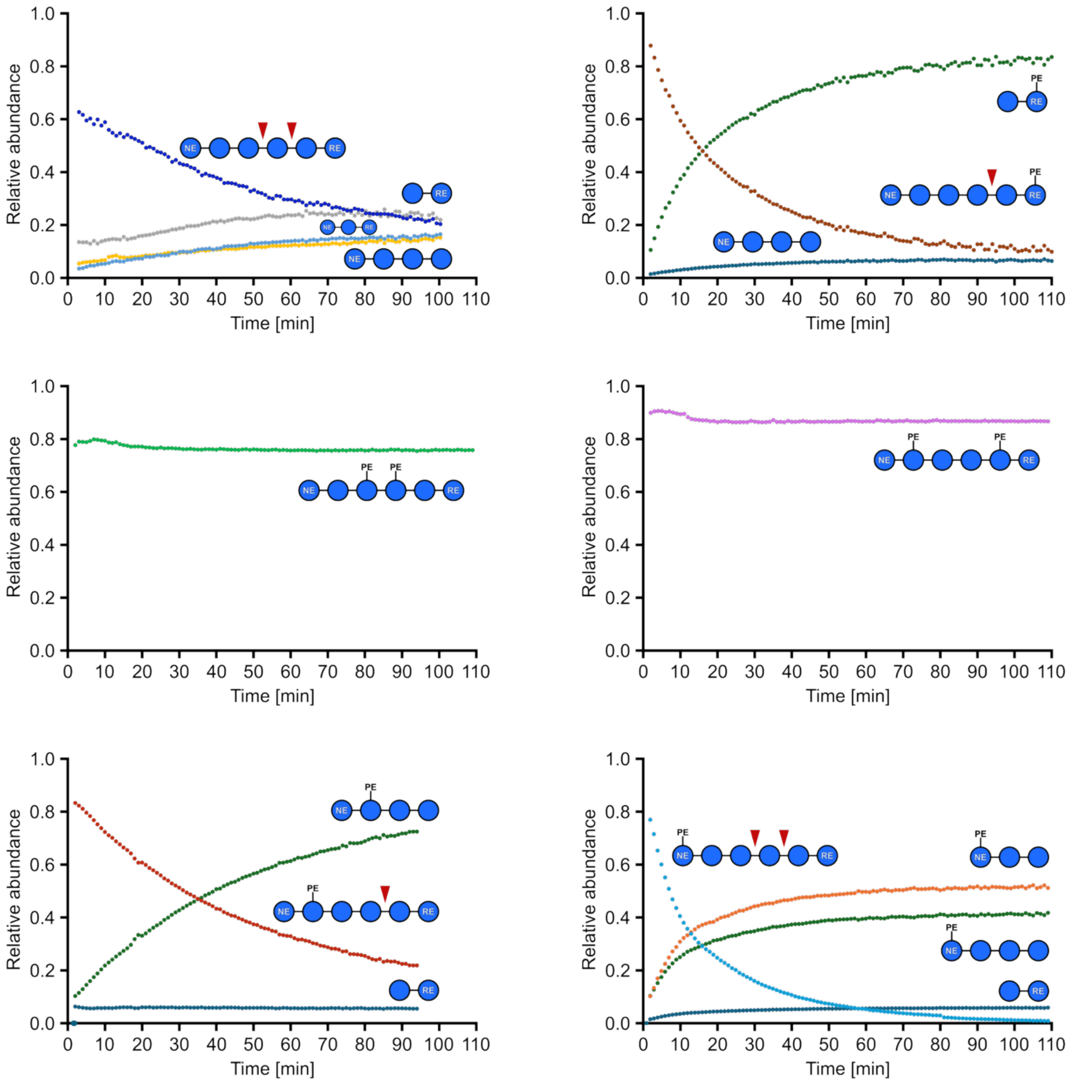


**Fig. S6.** Hydrolytic products detected by mass spectrometry for the indicated ligands.

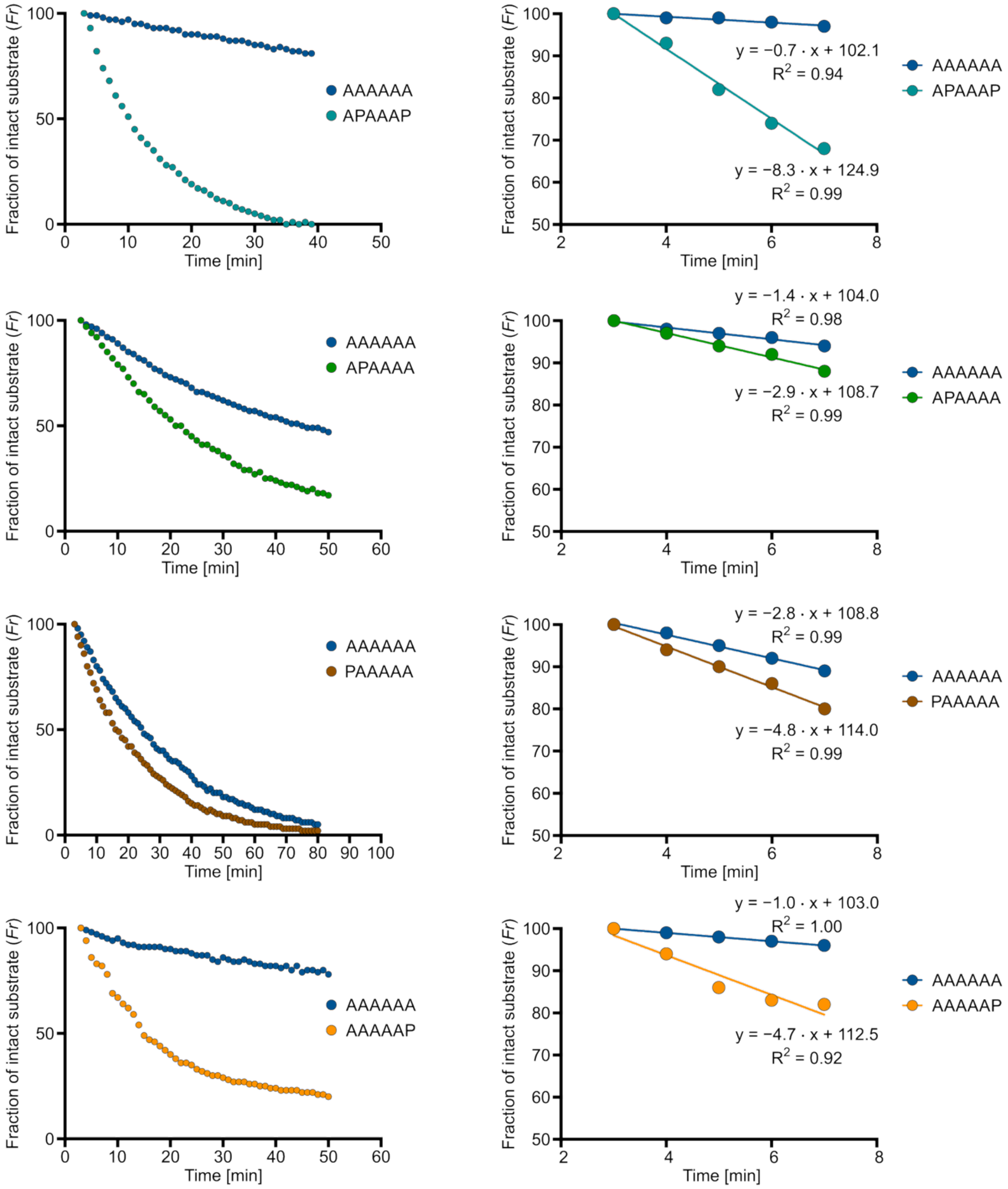


**Fig. S7.** Representative kinetics of ligand hydrolysis. Left column: MA detection of the indicated substrates in a reaction with wild type BcsZ. Right column: Linear fit of the ligand turnover in the indicated timeframe.

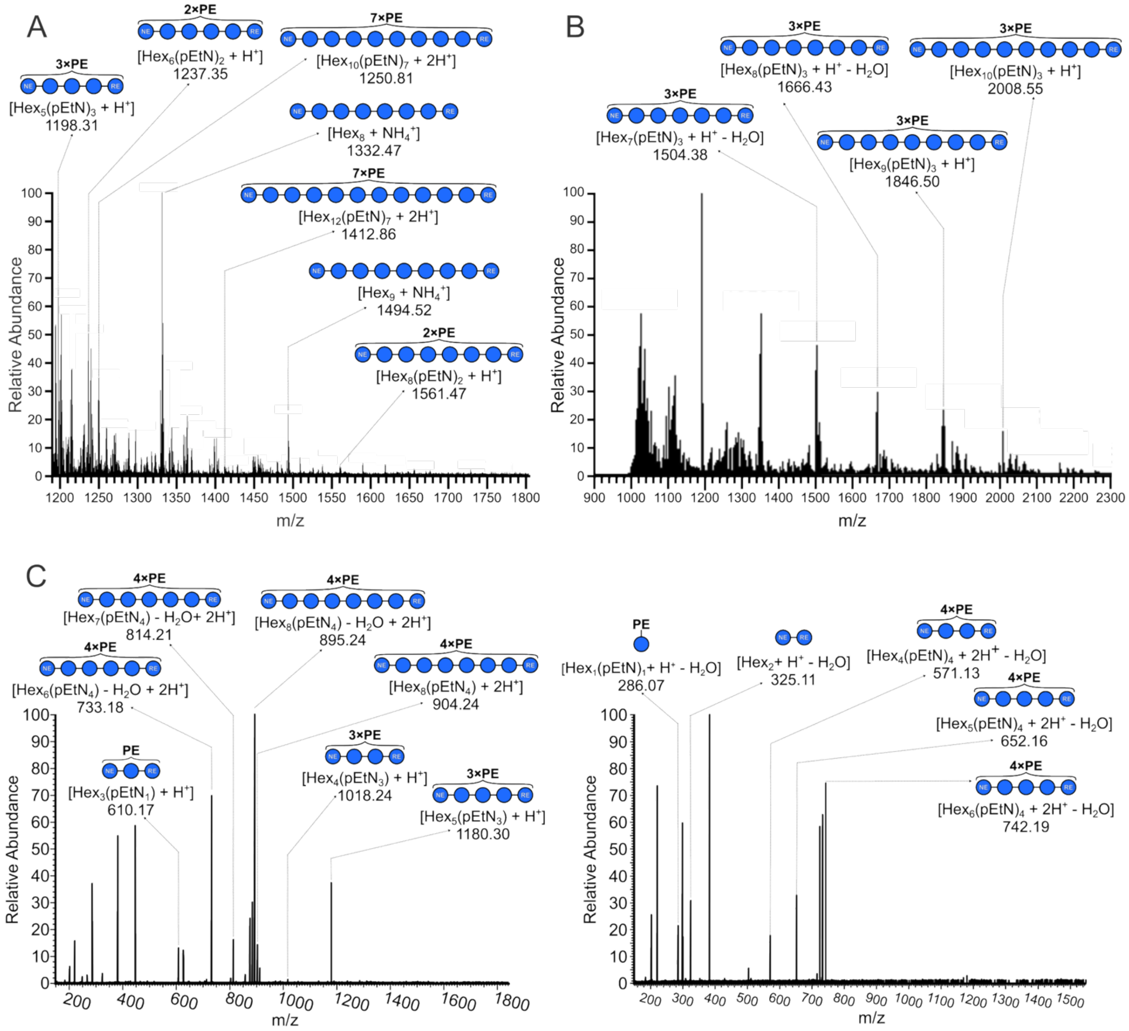


**Fig. S8.** Representative mass spectra of cello-oligosaccharides released from native pEtN cellulose digested with BcsZ. (A and B) Spectra obtained for samples derived from purified native pEtN cellulose (A) and digested pEtN cellulose-producing *E. coli* (B). (C) MS/MS fragmentation pattern of selected oligosaccharides.

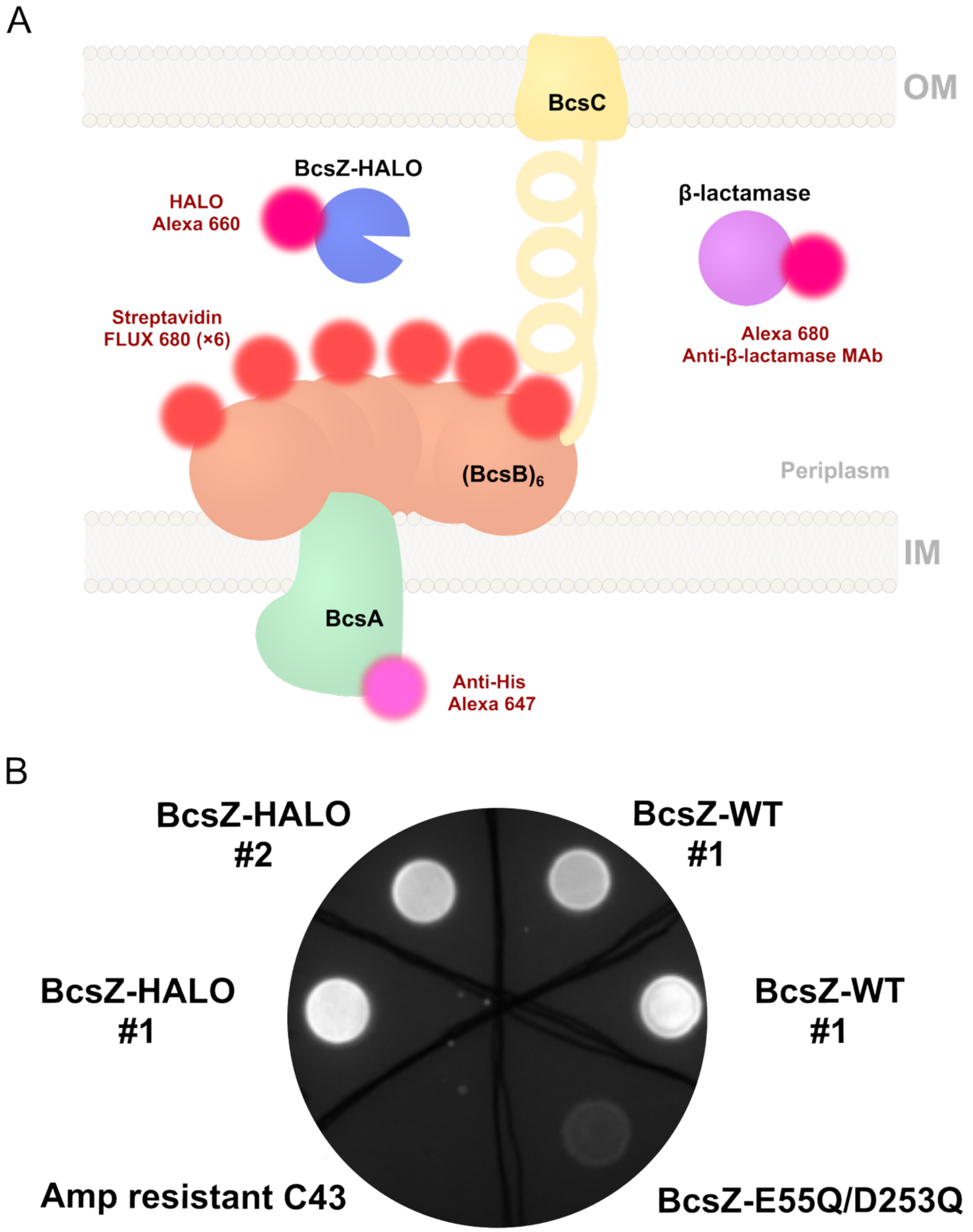


**Fig. S9.** *In vivo* BcsZ localization. (A) Cartoon illustration of the localization of labeled Bcs components. (B) Macrocolony staining with CongoRed as an indicator of pEtN cellulose production and secretion. The Halo-tagged BcsZ variant supports pEtN cellulose secretion.

v

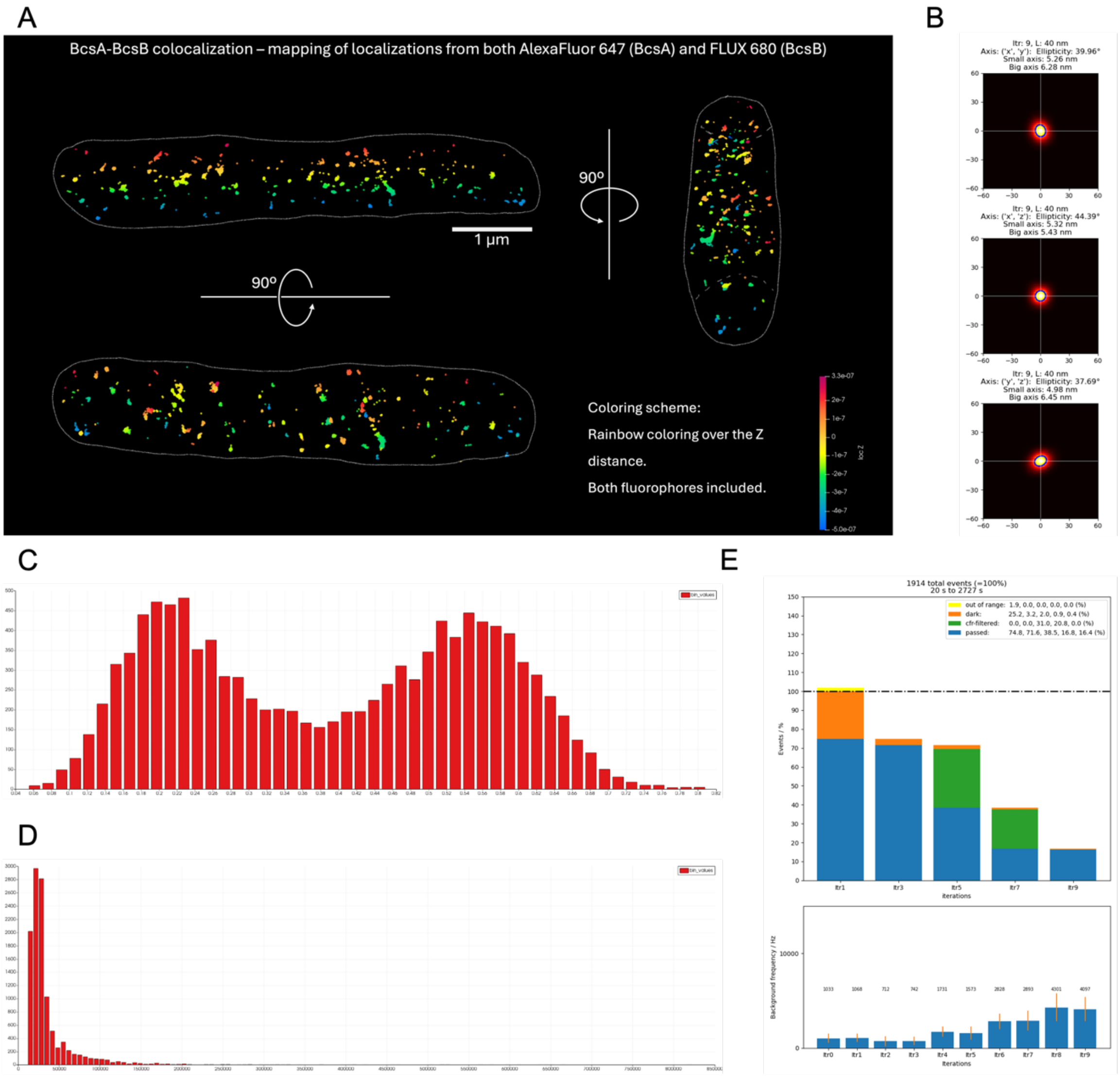


**Fig. S10.** MINFLUX analysis of BcsA and BcsB co-localization. (A) Representative 3D localization maps of BcsA labeled with anti-His Alexa Fluor 647 and BcsB labeled with streptavidin FLUX 680 are shown in three orthogonal views, with localization depth over the Z axis marked by rainbow coloring. (B) Corresponding localization precision maps,(C) DCR and (D) EFO distribution histograms, (E) fluorophore catching efficiency, and background intensity across ten MINFLUX iterations are shown. Cells were imaged in 16 mM MEA using 5 µg/mL of each labeling reagent.

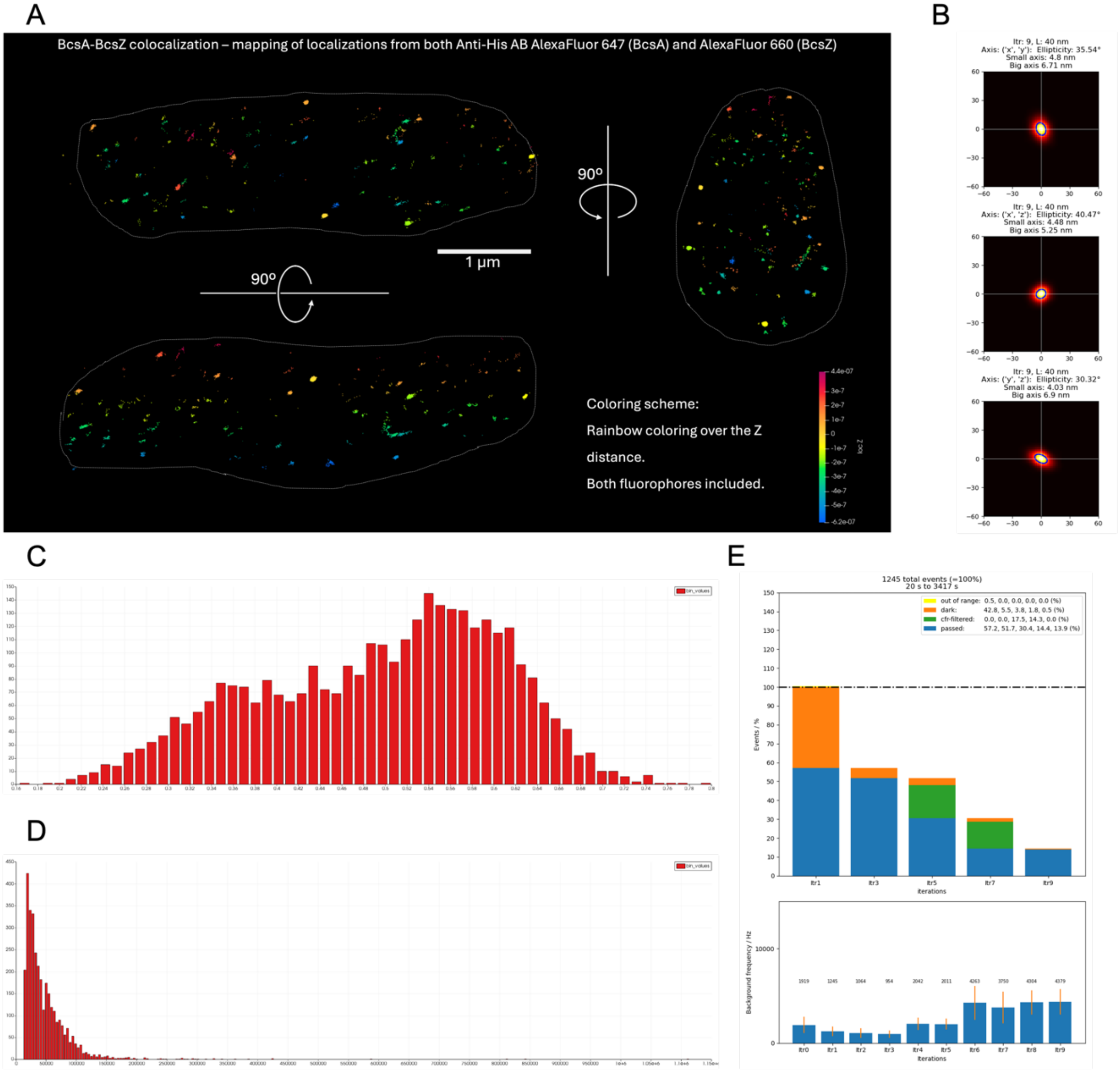


**Fig. S11**. MINFLUX analysis of BcsA and BcsZ co-localization. (A) Representative 3D localization maps of BcsA labeled with anti-His Alexa Fluor 647 and BcsZ labeled with HALO Alexa Fluor 660 are shown in three orthogonal views, with localization depth over the Z axis marked by rainbow coloring. (B) Corresponding localization precision maps, (C) DCR and (D) EFO distribution histograms, (E) fluorophore catching efficiency, and background intensity across ten MINFLUX iterations are shown. Cells were imaged in 16 mM MEA using 5 µg/mL of Anti-His antibodies and 200 nanomoles of HALO Alexa Fluor 660

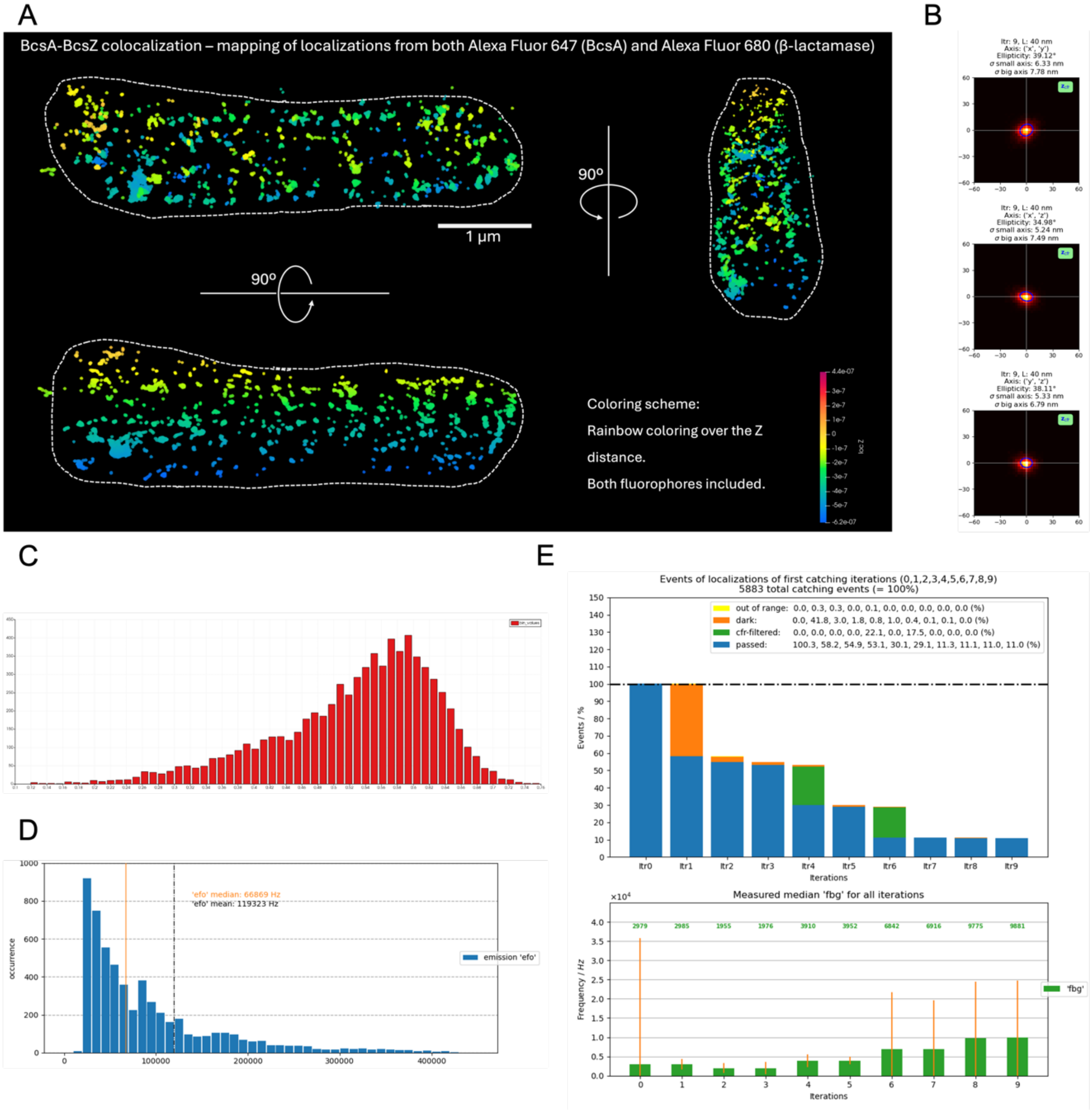


**Fig. S12.** MINFLUX analysis of BcsA and BcsZ co-localization. (A) Representative 3D localization maps of BcsA labeled with anti-His Alexa Fluor 647 and ß-lactamase labeled with anti- and ß-lactamase Alexa Fluor 680 are shown in three orthogonal views, with localization depth over the Z axis marked by rainbow coloring. Corresponding (B) localization precision maps, (C) DCR and (D) EFO distribution histograms, (E) fluorophore catching efficiency, and background intensity across ten MINFLUX iterations are shown. Cells were imaged in 16 mM MEA using 5 µg/mL of each labeling reagent.

Supplementary Table 1

| **Dataset** | **AAPPAA** | **APAAPA** | **APAAAP** | **APAAAA** |
|---|---|---|---|---|
| **Data collection** | | | | |
| Space group | P 1 21 1 | P 1 21 1 | P 1 21 1 | P 1 21 1 |
| Unit cell a, b, c (Å) | 90.61, 98.71, 93.30 | 88.70, 99.13, 91.66 | 90.62, 99.03, 93.18 | 90.50, 98.35, 92.94 |
| Unit cell α, β, γ (°) | 90.00, 103.47, 90.00 | 90.00, 101.84, 90.00 | 90.00, 103.10, 90.00 | 90.00, 102.87, 90.00 |
| Resolution range (Å) | 45.37–1.79 (1.82–1.79) | 49.30–1.69 (1.72–1.69) | 49.49–2.10 (2.14–2.10) | 49.42–1.90 (1.93–1.90) |
| Total observations | 1,041,716 (44,742) | 2,123,991 (83,535) | 644,188 (24,478) | 872,652 (40,019) |
| Unique reflections | 149,578 (6,970) | 172,651 (7,920) | 93,128 (4,144) | 122,439 (5,907) |
| Multiplicity | 7.0 (6.4) | 12.3 (10.5) | 6.9 (5.9) | 7.1 (6.8) |
| Completeness (%) | 99.7 (94.1) | 99.5 (92.4) | 99.5 (90.8) | 98.3 (95.2) |
| ⟨I/σ(I)⟩ | 7.7 (1.2) | 11.5 (1.4) | 11.0 (2.1) | 12.0 (1.9) |
| CC1/2 | 0.994 (0.240) | 0.999 (0.408) | 0.998 (0.740) | 0.998 (0.622) |
| $R_{merge}$ | 0.203 (1.787) | 0.149 (2.023) | 0.113 (0.727) | 0.117 (0.998) |
| $R_{meas}$ | 0.239 (2.125) | 0.163 (2.243) | 0.133 (0.871) | 0.137 (1.178) |
| $R_{pim}$ | 0.126 (1.137) | 0.065 (0.952) | 0.070 (0.475) | 0.072 (0.621) |

| **Dataset** | **PAAAAA** | **AAAAAP** | **AAAAAA** | **AAAAA** |
|---|---|---|---|---|
| **Data collection** | | | | |
| Space group | P 1 21 1 | P 1 21 1 | P1 | P1 |
| Unit cell a, b, c (Å) | 90.46, 98.12, 93.02 | 90.48, 99.12, 93.25 | 55.21, 86.91, 90.69 | 52.08, 84.61, 89.79 |
| Unit cell α, β, γ (°) | 90.00, 102.91, 90.00 | 90.00, 102.97, 90.00 | 71.34, 82.78, 78.9 | 72.52, 83.63, 81.26 |
| Resolution range (Å) | 49.39–2.00 (2.04–2.00) | 49.56–2.20 (2.25–2.20) | 41.98-1.40 (1.42-1.40) | 34.96/2.04 (2.08-2.04) |
| Total observations | 740,788 (34,281) | 554,439 (24,975) | 1030112 (49457) | 268690 (15231) |
| Unique reflections | 104,975 (5,074) | 80,582 (4,224) | 282109 (13302) | 83371 (4728) |

| | | | | |
|---|---|---|---|---|
| Multiplicity | 7.1 (6.8) | 6.9 (5.9) | 3.7 (3.7) | 3.2 (3.2) |
| Completeness (%) | 98.7 (96.7) | 99.5 (92.3) | 91.6 (87.1) | 90.4 (92.6) |
| ⟨I/σ(I)⟩ | 9.1 (1.7) | 9.2 (2.1) | 10.9 (1.2) | 9.3 (3.9) |
| CC1/2 | 0.996 (0.550) | 0.995 (0.604) | 0.998 (0.483) | 0.973 (0.702) |
| $R_{merge}$ | 0.169 (1.088) | 0.155 (0.796) | 0.059 (0.882) | 0.149 (0.43) |
| $R_{meas}$ | 0.199 (1.289) | 0.184 (0.961) | 0.069 (1.304) | 0.204 (0.578) |
| $R_{pim}$ | 0.104 (0.685) | 0.098 (0.532) | 0.035 (0.661) | 0.144 (0.408) |

| **Refinement** | **AAPPAA** | **APAAPA** | **APAAAP** | **APAAAA** |
|---|---|---|---|---|
| Resolution range used in refinement (Å) | 37.29–1.90 | 44.86–1.80 | 45.38–2.20 | 49.42–2.00 |
| Total reflections used in refinement | 125,623 | 143,231 | 81,327 | 105,442 |
| Rfree test set count | 6,542 | 7,248 | 4,008 | 5,186 |
| Rwork / Rfree | 0.1589 / 0.1916 | 0.1547 / 0.1924 | 0.1503 / 0.1850 | 0.1583 / 0.2079 |
| Number of non-H atoms | 13,019 | 13,096 | 12,566 | 13,010 |
| Protein atoms | 11,159 | 11,157 | 11,177 | 11,108 |
| Ligand/other atoms | 330 | 316 | 378 | 387 |
| Water atoms | 1,530 | 1,623 | 1,011 | 1,515 |
| Average B-factor, overall (Å²) | 30.59 | 30.19 | 40.53 | 27.97 |
| Average B-factor, protein (Å²) | 28.96 | 28.45 | 40.02 | 26.68 |
| Average B-factor, water (Å²) | 39.22 | 39.53 | 43.64 | 35.76 |
| Average B-factor, ligand/other (Å²) | 45.73 | 43.53 | 47.39 | 34.49 |
| RMSD bonds (Å) | 0.004 | 0.007 | 0.003 | 0.007 |
| RMSD angles (°) | 0.74 | 0.897 | 0.658 | 0.919 |
| Ramachandran favored / allowed / outliers (%) | 98.00 / 2.00 / 0.00 | 98.15 / 1.85 / 0.00 | 98.15 / 1.85 / 0.00 | 98.29 / 1.71 / 0.00 |
| Rotamer outliers (%) | 0.51 | 0.68 | 0.6 | 0.69 |

| | | | | |
|---|---|---|---|---|
| Clashscore | 4.03 | 3.24 | 2.92 | 3.57 |

| Refinement | PAAAAA | AAAAAP | AAAAAA | AAAAA |
|---|---|---|---|---|
| Resolution range used in refinement (Å) | 49.39–2.10 | 44.68–2.30 | 35.33-1.4 | 34.96-2.04 |
| Total reflections used in refinement | 91,274 | 71,320 | 281,888 | 83,332 |
| Rfree test set count | 4,649 | 3,552 | 14,217 | 4,198 |
| Rwork / Rfree | 0.1549 / 0.2140 | 0.1856 / 0.2435 | 0.1844/0.2103 | 0.1497/0.1988 |
| Number of non-H atoms | 13,153 | 12,527 | 13,038 | 12,453 |
| Protein atoms | 11,124 | 11,089 | 11,095 | 10,944 |
| Ligand/other atoms | 524 | 302 | 247 | 224 |
| Water atoms | 1,505 | 1,136 | 1,696 | 1,285 |
| Average B-factor, overall (Å$^2$) | 31.86 | 28.25 | 24.35 | 13 |
| Average B-factor, protein (Å$^2$) | 30.9 | 28.05 | 22.52 | 12 |
| Average B-factor, water (Å$^2$) | 38.62 | 29.91 | 34.52 | 20 |
| Average B-factor, ligand/other (Å$^2$) | 32.99 | 29.12 | 36.55 | 16 |
| RMSD bonds (Å) | 0.006 | 0.008 | 0.006 | 0.006 |
| RMSD angles (°) | 0.925 | 0.976 | 0.86 | 0.863 |
| Ramachandran favored / allowed / outliers (%) | 97.92 / 2.08 / 0.00 | 97.10 / 2.90 / 0.00 | 98.44/4.42/0 | 97.99/2.01/0 |
| Rotamer outliers (%) | 0.43 | 1.38 | 0.35 | 0.18 |
| Clashscore | 4.21 | 7.5 | 1.83 | 5.37 |

Supplementary Table 2:

| Classification | Composition | DP | pEtN groups | pEtN/Hex | Observed ion(s), m/z | Source(s) |
|---|---|---|---|---|---|---|
| Unmodified | Hex$_2$ | 2 | 0 | 0.00 | 343.12 $[M+H]^+$; 365.10 $[M+Na]^+$ | Cells; purified/extracted cellulose |
| | Hex$_3$ | 3 | 0 | 0.00 | 505.18 $[M+H]^+$; 527.15 $[M+Na]^+$ | Cells; purified/extracted cellulose |
| | Hex$_4$ | 4 | 0 | 0.00 | 667.23 $[M+H]^+$; 684.26 $[M+NH_4]^+$; 689.20 $[M+Na]^+$ | Cells; purified/extracted cellulose |
| | Hex$_5$ | 5 | 0 | 0.00 | 829.28 $[M+H]^+$; 846.31 $[M+NH_4]^+$; 851.26 $[M+Na]^+$ | Purified/extracted cellulose |
| | Hex$_6$ | 6 | 0 | 0.00 | 991.33 $[M+H]^+$; 1008.36 $[M+NH_4]^+$; 1013.31 $[M+Na]^+$ | Purified/extracted cellulose |
| | Hex$_7$ | 7 | 0 | 0.00 | 1153.39 $[M+H]^+$; 1170.42 $[M+NH_4]^+$; 1176.37 $[M+Na]^+$ | Purified/extracted cellulose |
| | Hex$_8$ | 8 | 0 | 0.00 | 1332.47 $[M+NH_4]^+$; 1338.42 $[M+Na]^+$ | Purified/extracted cellulose |
| | Hex$_9$ | 9 | 0 | 0.00 | 1494.52 $[M+NH_4]^+$ | Purified/extracted cellulose |
| Sparsely modified (<50% pEtN) | Hex$_5$-(pEtN)$_1$ | 5 | 1 | 0.20 | 952.29 $[M+H]^+$ | Cells; purified/extracted cellulose |
| | Hex$_4$-(pEtN)$_1$ | 4 | 1 | 0.25 | 790.24 $[M+H]^+$ | Cells; purified/extracted cellulose |
| | Hex$_8$-(pEtN)$_2$ | 8 | 2 | 0.25 | 1561.46 $[M+H]^+$ | Purified/extracted cellulose |
| | Hex$_7$-(pEtN)$_2$ | 7 | 2 | 0.29 | 1399.41 $[M+H]^+$ | Purified/extracted cellulose |
| | Hex$_6$-(pEtN)$_2$ | 6 | 2 | 0.33 | 1237.34 $[M+H]^+$ | Purified/extracted cellulose |
| | Hex$_9$-(pEtN)$_3$ | 9 | 3 | 0.33 | 1846.51 $[M+H]^+$ | Cells |
| | Hex$_8$-(pEtN)$_3$ | 8 | 3 | 0.38 | 842.70 $[M+2H]^{2+}$; 1684.46 $[M+H]^+$ | Cells; purified/extracted cellulose |
| | Hex$_5$-(pEtN)$_2$ | 5 | 2 | 0.40 | 1075.29 $[M+H]^+$ | Cells; purified/extracted cellulose |
| | Hex$_7$-(pEtN)$_3$ | 7 | 3 | 0.43 | 1522.41 $[M+H]^+$ | Purified/extracted cellulose |
| | Hex$_9$-(pEtN)$_4$ | 9 | 4 | 0.44 | 985.26 $[M+2H]^{2+}$; 1969.52 $[M+H]^+$ | Purified/extracted cellulose |
| Heavily modified (≥50% pEtN) | Hex$_4$-(pEtN)$_2$ | 4 | 2 | 0.50 | 913.24 $[M+H]^+$ | Cells; purified/extracted cellulose |
| | Hex$_6$-(pEtN)$_3$ | 6 | 3 | 0.50 | 1360.36 $[M+H]^+$ | Purified/extracted cellulose |
| | Hex$_8$-(pEtN)$_4$ | 8 | 4 | 0.50 | 904.23 $[M+2H]^{2+}$; 1807.47 $[M+H]^+$ | Purified/extracted cellulose |
| | Hex$_7$-(pEtN)$_4$ | 7 | 4 | 0.57 | 823.20 $[M+2H]^{2+}$; 1645.41 $[M+H]^+$ | Cells; purified/extracted cellulose |
| | Hex$_5$-(pEtN)$_3$ | 5 | 3 | 0.60 | 1198.31 $[M+H]^+$ | Cells; purified/extracted cellulose |
| | Hex$_8$-(pEtN)$_5$ | 8 | 5 | 0.63 | 965.75 $[M+2H]^{2+}$; 1930.48 $[M+H]^+$ | Purified/extracted cellulose |
| | Hex$_6$-(pEtN)$_4$ | 6 | 4 | 0.67 | 742.20 $[M+2H]^{2+}$; 1483.37 $[M+H]^+$ | Cells; purified/extracted cellulose |
| | Hex$_7$-(pEtN)$_5$ | 7 | 5 | 0.71 | 884.72 $[M+2H]^{2+}$; 1768.43 $[M+H]^+$ | Purified/extracted cellulose |
| | Hex$_5$-(pEtN)$_4$ | 5 | 4 | 0.80 | 1321.31 $[M+H]^+$ | Cells |
